# Illuminating phage dark matter beyond the reach of structural alignments with PhageFlare

**DOI:** 10.64898/2026.10.28.762509

**Authors:** Chonghao Wang, Yu Xu, Zeming Li, Louxin Zhang, Lu Zhang

## Abstract

Many phage proteins remain functionally uncharacterized despite advances in structural-similarity-based annotation. We introduce PhageFlare, a deep learning framework for predicting phage protein functions lacking significant structural matches to annotated references. Across three structure-disjoint benchmarks, where no significant Foldseek match was detected between training and test proteins (E-value thresholds of **10*^−^*^3^**, **10*^−^*^4^**, and **10*^−^*^2^**), PhageFlare achieved macro-averaged ***F*_1_** scores of 0.871, 0.889, and 0.738 outperforming the strongest baseline (0.789, 0.814, and 0.732). Systematic evaluations demonstrated that protein language model selection critically influences performance, with ESM-C achieving stronger overall performance than ESM-2 and ProtBert. Embedding-guided case studies revealed local structural correspondences among proteins with limited global similarity. Applied to the efam and Metagenomic Gut Virus datasets, PhageFlare annotated 38.0% and 51.8% of proteins unresolved by Fold-seek, respectively. Furthermore, PhageFlare identified lysogenic potential in 1,866 prophage-designated genomes misclassified as lytic by BACPHLIP. These results show that PhageFlare complements structural-similarity-based annotation and expands functional discovery across viral dark matter.

## Introduction

Bacteriophages (phages), viruses that infect bacteria, are ubiquitous and highly abundant components of environmental and host-associated microbial communities [1–7]. Phage genes encode proteins involved in diverse stages of the viral life cycle, including host recognition and genome entry, genome replication, virion assembly, host manipulation, and, during lytic infection, host-cell lysis. However, the extreme sequence diversity and rapid evolution of phage gene repertoires limit sequence-alignment-based functional annotation across distant relatives. Although metagenomic sequencing has greatly expanded known viral sequence space [8–12], functional interpretation has not kept pace. Consequently, many predicted phage proteins remain designated as hypothetical proteins or assigned to uncharacterized families [13–16]. This widening gap between sequence discovery and functional annotation underpins phage functional dark matter.

Unraveling this functional dark matter is critical both to advance phage biology and to translate phage mechanisms into biotechnological applications [17]. Protein functional annotation provides the mechanistic basis for reconstructing phage infection strategies, interpreting interactions with bacterial defense systems and understanding how phages influence microbial evolution and ecosystem processes [18–20]. However, low-throughput experimental validation cannot keep pace with the rapid expansion of viral sequence databases. Therefore, scalable computational inference is indispensable. Computational paradigms have progressively evolved: early methods relied on pairwise sequence homology [21, 22], whereas profile hidden Markov models (pHMMs) improved sensitivity by capturing position-specific conservation across protein families [23, 24]. Most recently, breakthroughs in protein structure prediction [25] coupled with rapid structural search algorithms, such as Foldseek [26], have enabled structural-alignment-based annotation at scale. Because tertiary structure is evolutionarily more conserved than primary sequence [27], structural alignment can detect remote homologs inaccessible to sequence-alignment-based approaches [28]. This transition is exemplified by Phold [29], a phage-specific framework that integrates ProstT5-derived structural representations, Foldseek and a curated database of predicted phage structures, achieving better annotation sensitivity compared to traditional sequence-alignment-based approaches in its reported benchmarks.

Structural alignments, however, cannot fully resolve phage functional dark matter [29, 30], as many phage proteins lack significant structural matches to functionally characterized reference proteins. Notably, benchmark evaluations demonstrate that protein language models (PLMs) can identify remote homologs overlooked by structural-alignment-based tools in other fields [31, 32], indicating that PLM embeddings capture complementary evolutionary signals. These findings suggest that PLMs may bridge current functional annotation gaps for phage proteins that remain unresolved by structural alignment.

Several phage-specific models have incorporated PLMs. VPF-PLM [15] uses PLM embeddings to classify phage proteins into nine broad functional labels, whereas Empathi [33] defines finer-grained functional labels and applies embedding-based classifiers to predict phage protein functions. GOPhage [34] employs a protein language model, ESM-2 [35], with genomic-context information to predict Gene Ontology terms [36] for phage proteins. While these efforts demonstrate the utility of PLM embeddings, previous studies have not explicitly stratified predictive performance by the presence or absence of significant structural alignments. Consequently, the degree to which PLM-based models improve annotation specifically for phage proteins lacking such significant hits remains poorly quantified. Furthermore, the extent to which PLM selection influences predictive performance in this setting remains unresolved. Although previous work benchmarked PLMs for phage protein annotation [15], the relative performance of the widely used ESM-2 and ESM-C [37] on proteins lacking significant structural matches remains unknown. Given that task-specific evaluations (e.g., Enzyme Commission classification) have demonstrated superior performance for ESM-2 over ESM-C [38], whether these task-specific rankings extend to phage protein function annotation warrants direct evaluation. Systematically evaluating PLM-based models on phage proteins lacking significant structural matches, and determining the effect of PLM selection on predictive performance, is therefore important for assessing the potential of PLMs for phage protein function annotation.

Here, we introduce PhageFlare, a deep learning framework designed to extend phage protein function annotation beyond the reach of structural alignment. We evaluated PhageFlare against Empathi [33], DHR [32], and HMMER [23] across structure-disjoint benchmarks (where test proteins lack significant structural matches to training proteins; see Methods) with different *E*-value thresholds. Foldseek [26] was excluded from evaluation, as it yielded no qualifying matches on the structure-disjoint benchmark. On the structure-disjoint benchmark with an *E*-value threshold of 10*^−^*^3^, PhageFlare achieved a macro-*F*_1_ score of 0.871, substantially outperforming the strongest competitor (0.789). PhageFlare similarly attained the highest *F*_1_ scores in the other two structure-disjoint benchmarks with *E*-values of 10*^−^*^2^ and 10*^−^*^4^. Systematic evaluation of underlying PLM backbones revealed that backbone selection critically influenced predictive performance, with ESM-C outperforming both ESM-2 and ProtBert [39]. Furthermore, embedding-guided structural analyses identified local structural correspondences among selected proteins located proximally in PhageFlare’s latent embedding space. To assess its utility for large-scale discovery, we applied PhageFlare to the efam [40] and Metagenomic Gut Virus (MGV) [41] datasets, where PhageFlare assigned candidate functions to 38.0% and 51.8% of proteins left unannotated by Foldseek, respectively. Finally, we investigated whether the enhanced annotation sensitivity from PhageFlare could translate into phage lifestyle classification and help recover lysogenic phages misclassified by a specialized phage lifestyle prediction model, BACPHLIP [42]. Integrating PhageFlare annotations successfully recovered 1,866 prophage-designated MGV genomes that BACPHLIP had misclassified as lytic. In summary, these results establish PhageFlare as a powerful complement to structural-alignment-based tools, expanding functional inference into phage dark matter and advancing the interpretation of phage genomes.

## Results

### Overview of PhageFlare

PhageFlare is designed using two complementary biological principles (Fig. 1A). First, different residues can contribute differently to protein function; for example, residues forming catalytic sites or interaction interfaces may be particularly important. Conventional protein representation strategies in PLMs rely on mean pooling across residue embeddings, implicitly assuming equal contributions from all amino acids. To account for different contributions of residues to a function, we designed a function-conditioned residue pooling module that enables different weighting of these residues. Second, the modular organization of phage genomes creates dependencies among neighboring genes [43], such that the predicted functions of nearby proteins can inform annotation of the target protein.

**Fig. 1:**
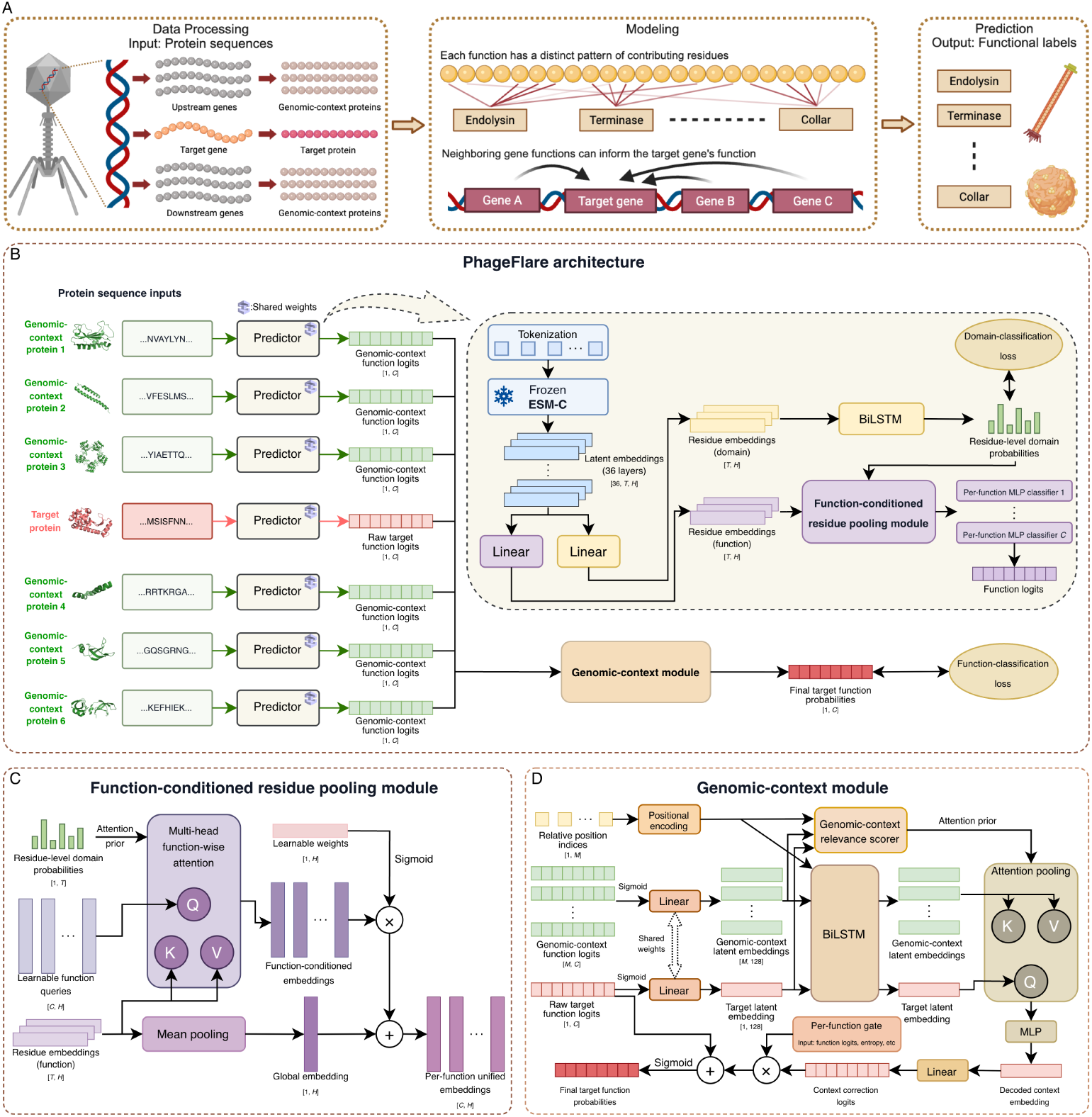
Overview of PhageFlare. (**A**) Conceptual framework. PhageFlare predicts phage protein functions by leveraging two biological insights: (i) distinct protein functions are associated with characteristic patterns of contributing amino acid residues, indicated by line intensity, and (ii) the functions of proteins encoded by neighboring genes can provide contextual information for predicting the function of a target protein. (**B**) PhageFlare architecture. The sequences of the target protein and its genomic-context proteins are processed by a shared-weight predictor. Each sequence is encoded by ESM-C to obtain latent embeddings of shape [36, *T* , *H*], where *T* denotes sequence length and *H* represents the hidden dimension. These embeddings are used to predict residue-level domain probabilities and protein function logits through a function-conditioned residue pooling module and per-function multilayer perceptron (MLP) classifiers (1 to *C*, where *C* represents the number of functional labels). The function logits of the target and genomic-context proteins are subsequently integrated by the genomic-context module to produce the final target-function probabilities. (**C**) Function-conditioned residue pooling module. Learnable function queries perform attention over residue embeddings, incorporating residue-level domain probabilities as an attention prior to derive function-conditioned embeddings. These embeddings are merged with a global embedding obtained by mean pooling to generate per-function unified embeddings. (**D**) Genomic-context module. This module refines raw target function logits by incorporating genomic context (*M* genomic-context proteins in total). A genomic-context relevance scorer serves as an attention prior that modulates the pooling weight assigned to each genomic-context protein. A per-function gate controls the extent to which genomic-context information contributes to the prediction of each function.

PhageFlare applies a shared sequence predictor to the target protein and its genomic-context proteins (Fig. 1B). The predictor uses a frozen, pretrained ESMC protein language model to obtain residue-level latent embeddings. Two layer-wise learned combinations of the ESM-C latent embeddings are projected to derive specialized embeddings for residue-level domain prediction and protein function prediction, respectively. PhageFlare employs function-conditioned residue pooling to identify a weighted summary of residue embeddings that are informative for each function (Fig. 1C). Specifically, a learnable query for each function attends to the residue embeddings through multi-head attention, thereby assigning function-specific weights to all residues. Predicted residue-level domain probabilities are incorporated as an attention prior, biasing the model towards residues that are more likely to carry functionally relevant information. Each resulting function-conditioned embedding is combined, through a learned feature-wise gate, with a global mean-pooled embedding of the protein. Separate MLP classifiers then convert these per-function unified embeddings into predicted function logits.

The genomic-context module subsequently calibrates the target protein predictions using the predictions for genomic-context proteins (Fig. 1D). The function probabilities of the target and genomic-context proteins are projected into a 128-dimensional latent space using a linear projection. Relative-position embeddings preserve the order and relative gene offset of each genomic-context protein. A bidirectional long short-term memory (BiLSTM) [44, 45] network encodes the ordered target–context representation. For each genomic-context protein, a relevance scorer integrates its latent embedding, the target latent embedding, relative position and the entropy from genomic-context function probabilities. This score serves as an attention prior that modulates the contribution of each genomic-context protein during attention pooling.

The decoded context embedding is turned into context correction logits through a linear projection. As genomic context may be informative for some functions but uninformative for others, PhageFlare uses a separate gate for each function to regulate the contribution of the context-derived correction. These gates are estimated from the target and context-correction logits, together with summary measures of prediction uncertainty and contextual relevance. The final target function logits are obtained by adding the gated context correction to the raw target function logits from the predictor. When no valid genomic-context proteins are available, the predictions from the predictor are retained unchanged. PhageFlare yields predictions for 45 functional labels (Extended Data Fig. 1). If no functional label is assigned to a target protein, it is assigned the ’unknown’ label. These 45 functional labels originate from Empathi [33].

### PhageFlare retains predictive performance in the absence of significant structural alignments

To evaluate whether PhageFlare can annotate phage proteins beyond the reach of structural alignment, we constructed structure-disjoint benchmarks in which no significant Foldseek match was detected between any training and test protein in either search direction across three *E*-value thresholds: *E ≤* 10*^−^*^3^ (Fig. 2A–C), *E ≤* 10*^−^*^4^ (Fig. 2D–F) and *E ≤* 10*^−^*^2^ (Fig. 2G–I; Methods). Because Foldseek returned no qualifying structural matches under these definitions, it could not generate predictions and was excluded from the evaluation. We compared PhageFlare against the PLM-based model Empathi, Empathi (tuned thresholds), the embedding-retrieval method Dense Homolog Retriever (DHR), as well as HMMER. Empathi was evaluated using two thresholding schemes: Empathi used the default threshold of 0.5 for all functions, whereas Empathi (tuned thresholds) used function-specific thresholds optimized on the validation set. In addition, to assess how the choice of pretrained PLM backbone influences functional prediction in the absence of structural homologs, we evaluated PhageFlare across three backbones: ESM-C, ESM-2 and ProtBert.

**Fig. 2:**
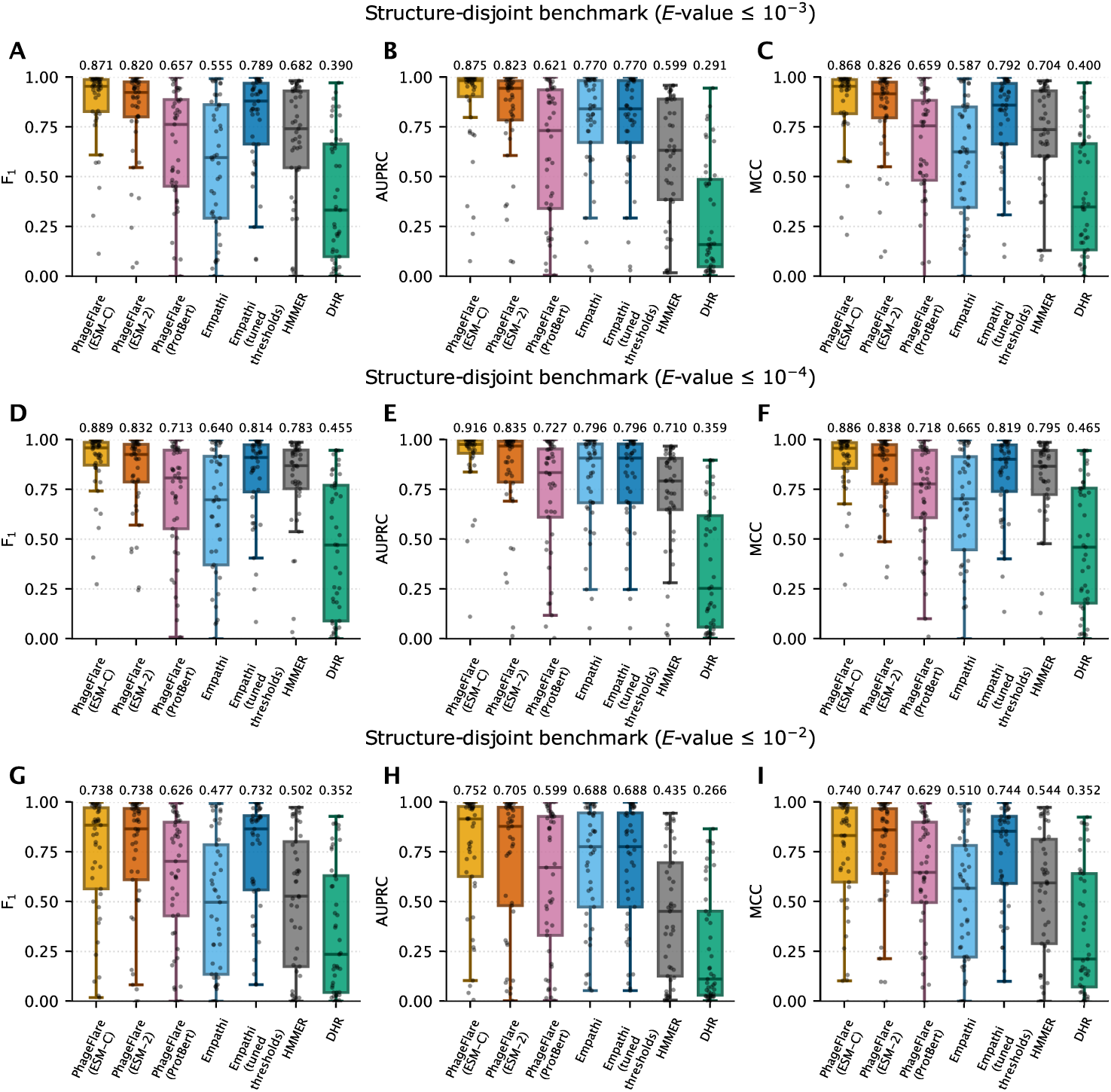
Comparative performance of PhageFlare on structure-disjoint benchmarks across different *E*-value thresholds. (**A–I**) Distributions of label-wise *F*_1_ score (**A,D,G**), AUPRC (**B,E,H**) and MCC (**C,F,I**) on structure-disjoint test datasets with Foldseek *E*-value thresholds of 10*^−^*^3^ (**A–C**), 10*^−^*^4^ (**D–F**) and 10*^−^*^2^ (**G–I**). Under each threshold, no significant Foldseek alignment was detected between training and test proteins in either search direction. PhageFlare variants using different pretrained PLM backbones (PhageFlare (ESM-C), PhageFlare (ESM-2) and PhageFlare (ProtBert)) were compared with Empathi, Empathi (tuned thresholds), HMMER and DHR. Empathi and Empathi (tuned thresholds) denote predictions obtained using a universal decision threshold of 0.5 and function-specific thresholds optimized on the validation dataset, respectively. Each point represents one of the 45 functional labels (*n* = 45), and values above the boxes indicate macro averages across all functional labels.

On the primary structure-disjoint benchmark (*E ≤* 10*^−^*^3^), PhageFlare (ESM-C) retained high predictive performance across the 45 functional labels, achieving macro-averaged *F*_1_, area under the precision–recall curve (AUPRC) and Matthews correlation coefficient (MCC) values of 0.871, 0.875 and 0.868, respectively (Fig. 2A–C). Paired protein-level bootstrap resampling across the test proteins confirmed narrow 95% confidence intervals (CIs) for both macro-*F*_1_ (0.867–0.874) and macro-MCC (0.864–0.871; Supplementary Fig. 1). Empathi (tuned thresholds) was the strongest external comparator, attaining macro-averaged *F*_1_, AUPRC and MCC values of 0.789 (95% CI: 0.783–0.793), 0.770 and 0.792 (95% CI: 0.787–0.797), respectively. PhageFlare (ESMC) outperformed Empathi (tuned thresholds) across all metrics by margins of 0.082 in *F*_1_, 0.105 in AUPRC and 0.076 in MCC. Although optimizing function-specific thresholds substantially improved the *F*_1_ and MCC of Empathi relative to its universal 0.5 threshold, a clear performance gap with PhageFlare remained. HMMER and DHR exhibited more pronounced reductions in performance: HMMER attained macro-averaged *F*_1_, AUPRC and MCC values of 0.682 (95% CI: 0.675–0.688), 0.599 and 0.704 (95% CI: 0.697–0.709), whereas DHR reached 0.390 (95% CI: 0.383–0.396), 0.291 and 0.400 (95% CI: 0.392–0.407), respectively. Across individual functional categories at *E ≤* 10*^−^*^3^, PhageFlare (ESM-C) achieved an *F*_1_ score of at least 0.90 for 31 of the 45 labels and exceeded Empathi (tuned thresholds) on 29 labels.

Evaluating the models at *E ≤* 10*^−^*^4^ and *E ≤* 10*^−^*^2^ further characterized predictive performance across varying degrees of structural divergence (Fig. 2D–I and Supplementary Fig. 1). Under the *E ≤* 10*^−^*^4^ threshold, which excludes closer structural matches while retaining more distant relationships, performance increased across all methods (Fig. 2D–F). PhageFlare (ESM-C) achieved the highest macro-averaged *F*_1_, AUPRC and MCC values of 0.889 (95% CI: 0.885–0.893), 0.916 and 0.886 (95% CI: 0.882–0.890), exceeding Empathi (tuned thresholds) (0.814, 0.796 and 0.819), HMMER (0.783, 0.710 and 0.795) and DHR (0.455, 0.359 and 0.465). Conversely, applying the *E ≤* 10*^−^*^2^ threshold reduced performance across all methods, with HMMER showing the steepest declines in macro-*F*_1_, to 0.502 (Fig. 2G–I). Under this most restrictive regime, PhageFlare (ESM-C) maintained a macro-*F*_1_ of 0.738 (95% CI: 0.732–0.743), a macro-MCC of 0.740 (95% CI: 0.734–0.745) and the highest macro-AUPRC of 0.752, outperforming Empathi (tuned thresholds) in AUPRC (0.688) while achieving comparable *F*_1_ (0.732) and MCC (0.744).

Comparing the three PhageFlare variants demonstrated that PLM backbone selection strongly influences phage protein function prediction (Fig. 2A–I and Supplementary Fig. 1). Across all three *E*-value thresholds, both ESM-family backbones (ESM-C and ESM-2) substantially outperformed ProtBert, which attained macro-*F*_1_ scores of 0.657, 0.713 and 0.626 at *E ≤* 10*^−^*^3^, 10*^−^*^4^ and 10*^−^*^2^, respectively. When equipped with the ESM-2 backbone, the same pretrained PLM used by Empathi, PhageFlare (ESM-2) still outperformed Empathi (tuned thresholds) at both *E ≤* 10*^−^*^3^ (*F*_1_ = 0.820 versus 0.789; AUPRC = 0.823 versus 0.770; MCC = 0.826 versus 0.792) and *E ≤* 10*^−^*^4^ (*F*_1_ = 0.832 versus 0.814; AUPRC = 0.835 versus 0.796; MCC = 0.838 versus 0.819). Changing the backbone from ESM-2 to ESM-C further increased macro-averaged *F*_1_ and AUPRC by 0.051 and 0.052 at *E ≤* 10*^−^*^3^, and by 0.057 and 0.081 at *E ≤* 10*^−^*^4^. At *E ≤* 10*^−^*^2^, ESM-C and ESM-2 achieved identical macro-*F*_1_ (0.738) and similar macro-MCC (0.740 versus 0.747), while ESM-C retained a substantial lead in macro-AUPRC (0.752 versus 0.705). Consequently, ESM-C was selected as the default PLM backbone for PhageFlare, and used in all subsequent experiments. In summary, these findings establish that PhageFlare retains strong and functionally broad predictive performance in the absence of significant structural alignments, with the ESM-C backbone providing the most effective representations for resolving phage functional dark matter.

### Structural characterization of local relationships in PhageFlare’s embedding space

PhageFlare’s retained performance on the structure-disjoint benchmark prompted us to examine the relationships between training and test proteins in its latent embedding space. We analyzed target protein embeddings from the portal and major capsid classifiers, extracted before genomic-context integration (Methods). By using t-distributed stochastic neighbor embedding (t-SNE) projections [46], training and test proteins carrying the same functional label showed pronounced separation, although some local neighborhoods contained samples from both datasets (Fig. 3A and Extended Data Fig. 2A). In contrast, the corresponding projections in the sequence-disjoint setting, where test proteins share no sequence clusters with training proteins (Methods), demonstrated many overlapping neighborhoods (Extended Data Fig. 3). This divergence indicates that underlying structural relatedness shapes the geometry of PhageFlare’s latent embeddings.

**Fig. 3:**
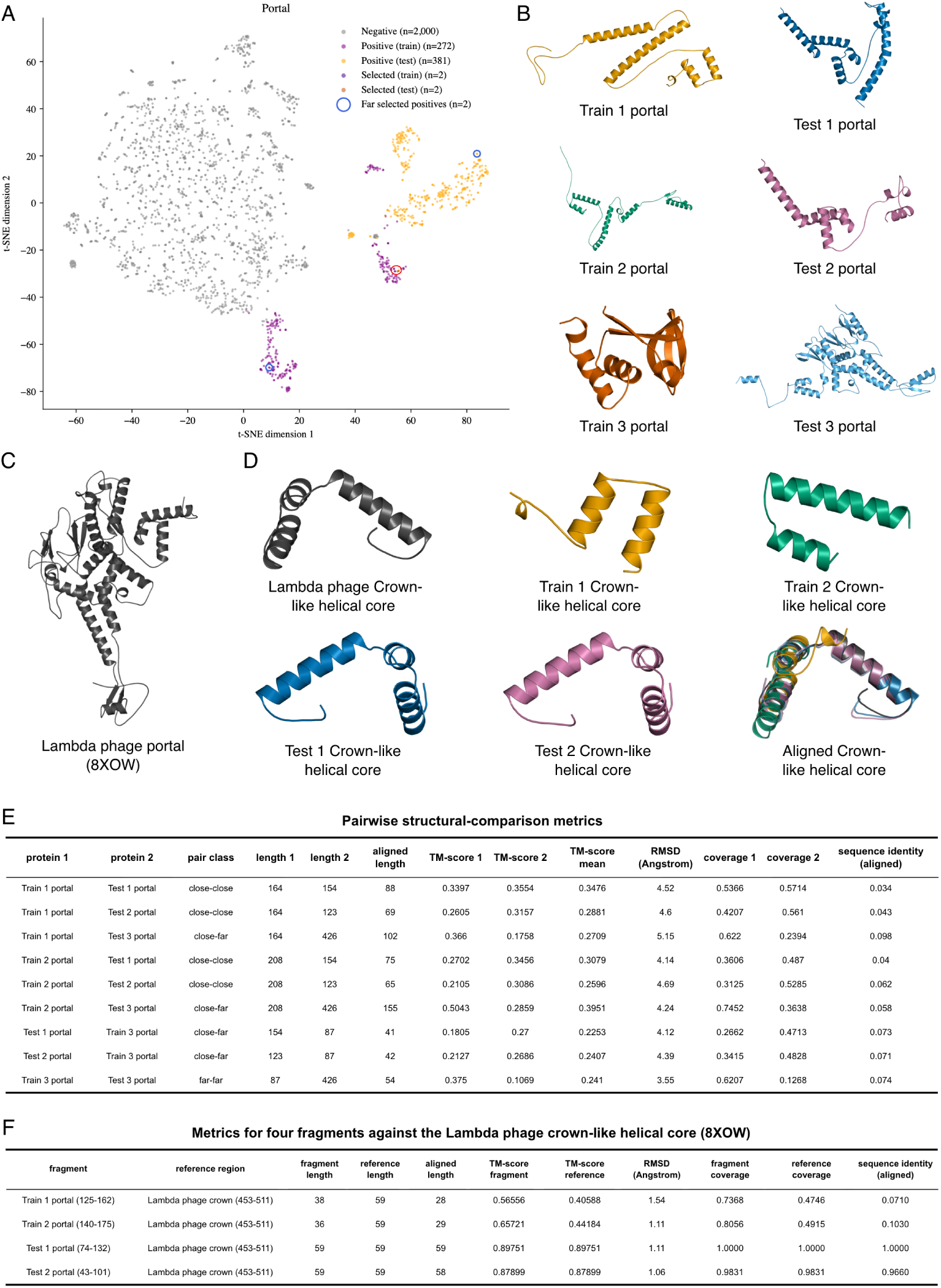
Embedding-guided analysis of portal proteins in the structure-disjoint benchmark. (**A**) Two-dimensional t-SNE projection of embeddings extracted from the PhageFlare portal classifier, comprising 272 training positives, 381 test positives and 2,000 sampled negatives lacking the portal label. Four proximal samples (Train 1, Train 2, Test 1 and Test 2) and two more distant samples (Train 3 and Test 3) were used for structural analysis. (**B**) AlphaFold 3 predicted single-chain structures of the six representative portal proteins. (**C**) Experimentally determined Lambda phage portal subunit used as the structural reference (PDB 8XOW, portal protein B subunit). (**D**) Crown-like helical regions extracted from the four embedding-proximal proteins and the reference Lambda portal crown. The final subpanel shows their US-align-derived superposition onto the reference. (**E**) Metrics for all nine comparisons between the three training and three test samples. Pair classes indicate selection from proximal or distant positions in the t-SNE projection. (**F**) Metrics for comparisons of the four crown-like fragments with the reference crown.

To investigate the biological context of the localized overlap, we analyzed six representative portal proteins from the structure-disjoint benchmark. These comprised a tightly clustered group of four samples (two training and two test) within the t-SNE projection, alongside two samples (one training and one test) located farther from this group (Fig. 3A,B). Across the four train–test comparisons within the proximal group, US-align [47] yielded mean TM-scores of 0.260–0.348, root-mean-square deviation (RMSD) of 4.14–4.69° A and aligned sequence identities of 3.4–6.2% (Fig. 3E). Thus, proximity in the projection did not require high overall similarity between these proteins.

Despite their limited overall similarity, the four proximal samples contained helical regions with partial structural correspondence to the crown-like helical core of the Lambda phage portal protein (PDB 8XOW [48]; Fig. 3C,D). The crown-like helical core comprises residues 453–511 of gpB [49]. The selected fragments from the two training proteins aligned to this reference over 28 and 29 residues, with fragment-normalized TM-scores of 0.57 and 0.66, and RMSDs of 1.54 and 1.11° A, respectively (Fig. 3F). These alignments covered approximately half of the reference crown, supporting a local correspondence. Furthermore, the corresponding regions in the two test proteins exhibited an even higher structural concordance with the Lambda crown, representing closely related portal fragments.

A second case study examined major capsid proteins within a distinct localized train–test neighborhood (Extended Data Fig. 2). The selected training protein and two test proteins had mean train–test TM-scores of 0.371 and 0.408. Their local helical regions were compared with the P-domain spine helix of HK97 gp5, defined by residues 200–239 [50] (PDB 1OHG [51]). The regions from Train 1 and Test 1 each aligned to the reference over 29 residues, with RMSDs of 0.77 and 0.96° A, respectively, despite an aligned sequence identity of 3.4% in both comparisons. Test 2 contained a more strongly bent region and showed a more limited correspondence, with 23 aligned residues and an RMSD of 2.69° A.

In summary, these analyses demonstrate that proximity in PhageFlare’s latent embedding space can reflect conserved local structural motifs even in the absence of global structural similarity. These observations provide a structural context for latent embedding proximity in the absence of significant Foldseek matches to the training set.

### Functional discrimination in classifier embeddings and attention profiles of PhageFlare

To characterize the organization of PhageFlare’s learned embedding space, we assessed whether its function-specific classifier embeddings could distinguish positive from negative samples in the structure-disjoint test dataset with the *E*-value threshold of 10*^−^*^3^. These embeddings were extracted prior to genomic-context integration and visualized independently for each functional label using t-SNE (Methods). As shown in Fig. 4A–F, positive samples were concentrated in characteristic regions of the projections for phage virion proteins, major capsid, DNA polymerase, terminase, anti-restriction and reductase. Furthermore, projections for the remaining 39 functions showed analogous patterns (Supplementary Figs. 2–6). These observations provide qualitative evidence that PhageFlare’s classifier embeddings adopt a highly function-discriminative topology, effectively segregating unseen test proteins.

**Fig. 4:**
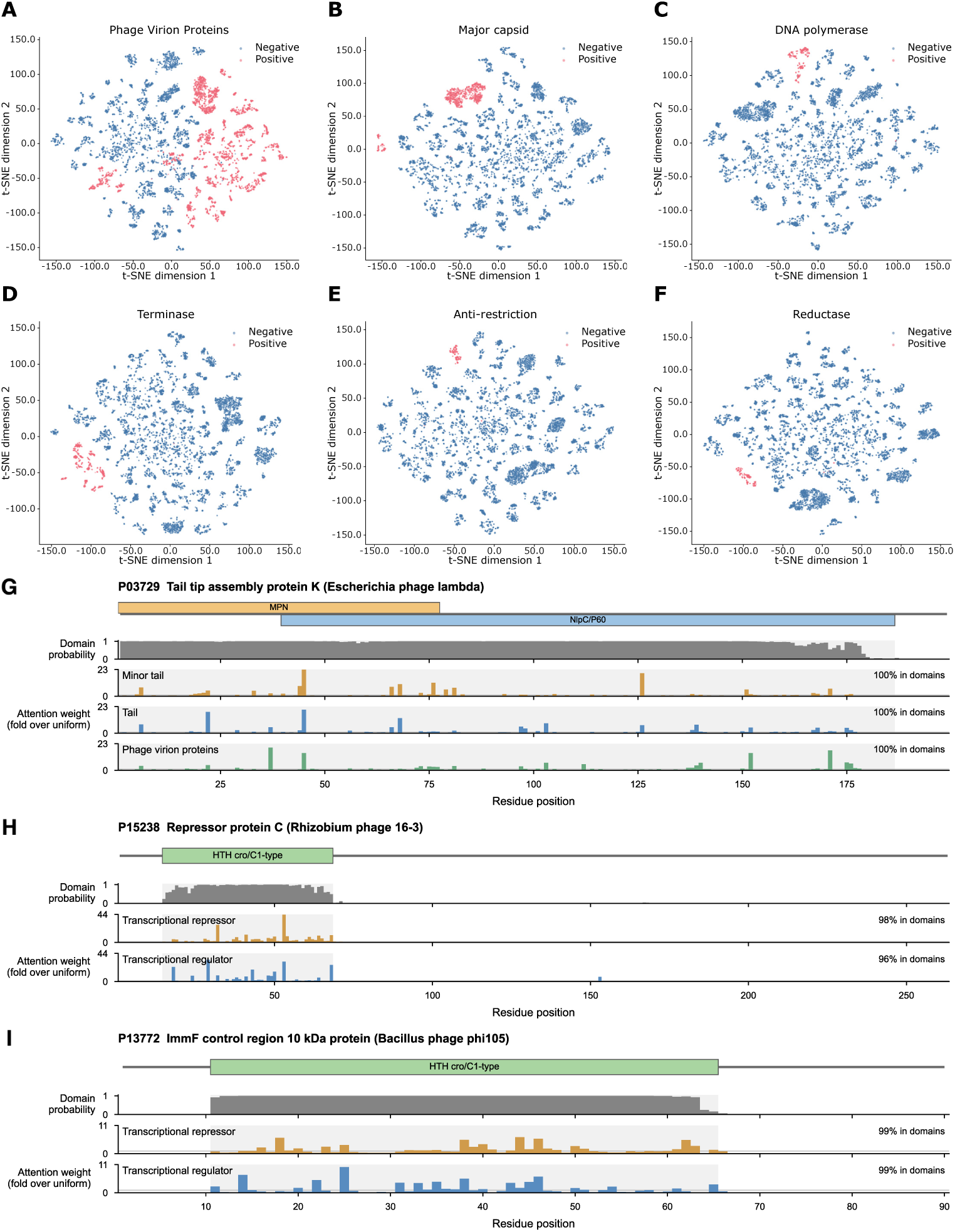
Function-specific classifier embeddings and residue-level attention analysis. (**A**–**F**) Two-dimensional t-SNE projections of target protein embeddings from the PhageFlare classifiers for phage virion proteins (**A**), major capsid (**B**), DNA polymerase (**C**), terminase (**D**), anti-restriction (**E**) and reductase (**F**) in the structure-disjoint test dataset (*E*-value *≤* 10*^−^*^3^). (**G**–**I**) Attention profiles with annotated domains and predicted domain probabilities for three representative phage proteins: tail tip assembly protein K (**G**), repressor protein C (**H**), and ImmF control region 10 kDa protein (**I**). Colored rectangles indicate UniProt-annotated domains, and gray shading marks corresponding domain regions. Attention tracks display residue-level attention weights for relevant functional labels, normalized as fold enrichment relative to uniform attention.

To examine how PhageFlare derives these function-conditioned representations at the residue level, we analyzed the predicted domain probabilities and head-averaged attention weights for three representative phage proteins evaluated without genomic context (Fig. 4G–I). None of the three proteins was present in the function and domain training datasets or had a significant MMseqs2 alignment to them (*E*-value *≤* 10*^−^*^3^). Across all three proteins, PhageFlare’s predicted residue-level domain probabilities broadly corresponded to the UniProt-annotated domain regions. In the tail tip assembly protein K (UniProt P03729), high predicted domain probabilities spanned the overlapping N-terminal MPN domain and C-terminal NlpC/P60 domain (Fig. 4G). Similarly, predicted domain probabilities were tightly confined to the N-terminal HTH cro/C1-type domain in the repressor protein C (P15238; Fig. 4H) and to the central HTH cro/C1-type domain in the ImmF control region 10 kDa protein (P13772; Fig. 4I).

Guided by the predicted domain prior, the model concentrated nearly all residue-level attention mass within the annotated domain regions. For tail tip assembly protein K (P03729), 100% of the attention mass for its ground-truth functional queries—minor tail, tail and phage virion proteins—localized within the MPN and NlpC/P60 domains (Fig. 4G). For the two transcriptional repressors, the HTH cro/C1-type domain captured 98% and 96% of the attention mass for the transcriptional repressor and transcriptional regulator queries in P15238 (Fig. 4H), and 99% for both queries in P13772 (Fig. 4I). Whereas the overall in-domain attention share was governed primarily by the shared domain prior, individual function queries attended to distinct, sparse subsets of residues within each domain, yielding query-specific enrichment peaks of up to approximately 23-fold, 44-fold and 11-fold over uniform attention in P03729, P15238 and P13772, respectively. Collectively, these profiles illustrate how domain-informed attention pooling restricts aggregation to functionally relevant domains while enabling individual function queries to weight distinct residue positions within them.

### PhageFlare expands functional annotation coverage across environmental phage proteomes

To determine whether the performance observed on the structure-disjoint benchmark could translate into broader annotation coverage in large phage protein collections, we applied PhageFlare to the dereplicated and filtered efam-XC protein set (efam) and proteins from complete genomes in the MGV dataset. The efam dataset comprised proteins without genomic context, whereas the MGV dataset included target proteins with their genomic-context proteins (Methods). Foldseek served as the primary baseline to evaluate annotation expansion beyond significant structural alignments, whereas Empathi, VPF-PLM, HMMER and MMseqs2 provided complementary benchmarks for PLM-based and sequence-based prediction approaches, respectively. Annotations were assigned using the method-specific decision thresholds and search criteria described in Methods.

PhageFlare assigned functional annotations to 70.1% of the efam proteins, compared with 62.2% for Foldseek (Fig. 5A). This coverage expansion was even more pronounced in MGV: PhageFlare annotated 71.3% of the total proteins, compared with 55.2% for Foldseek, marking a relative increase of 29.1% (Fig. 5B). PhageFlare likewise outperformed all other baselines in overall coverage: Empathi annotated 61.1% (efam) and 45.9% (MGV) of proteins, HMMER annotated 49.8% and 33.2%, MMseqs2 annotated 47.4% and 31.1%, and VPF-PLM annotated 37.5% and 36.4%.

**Fig. 5:**
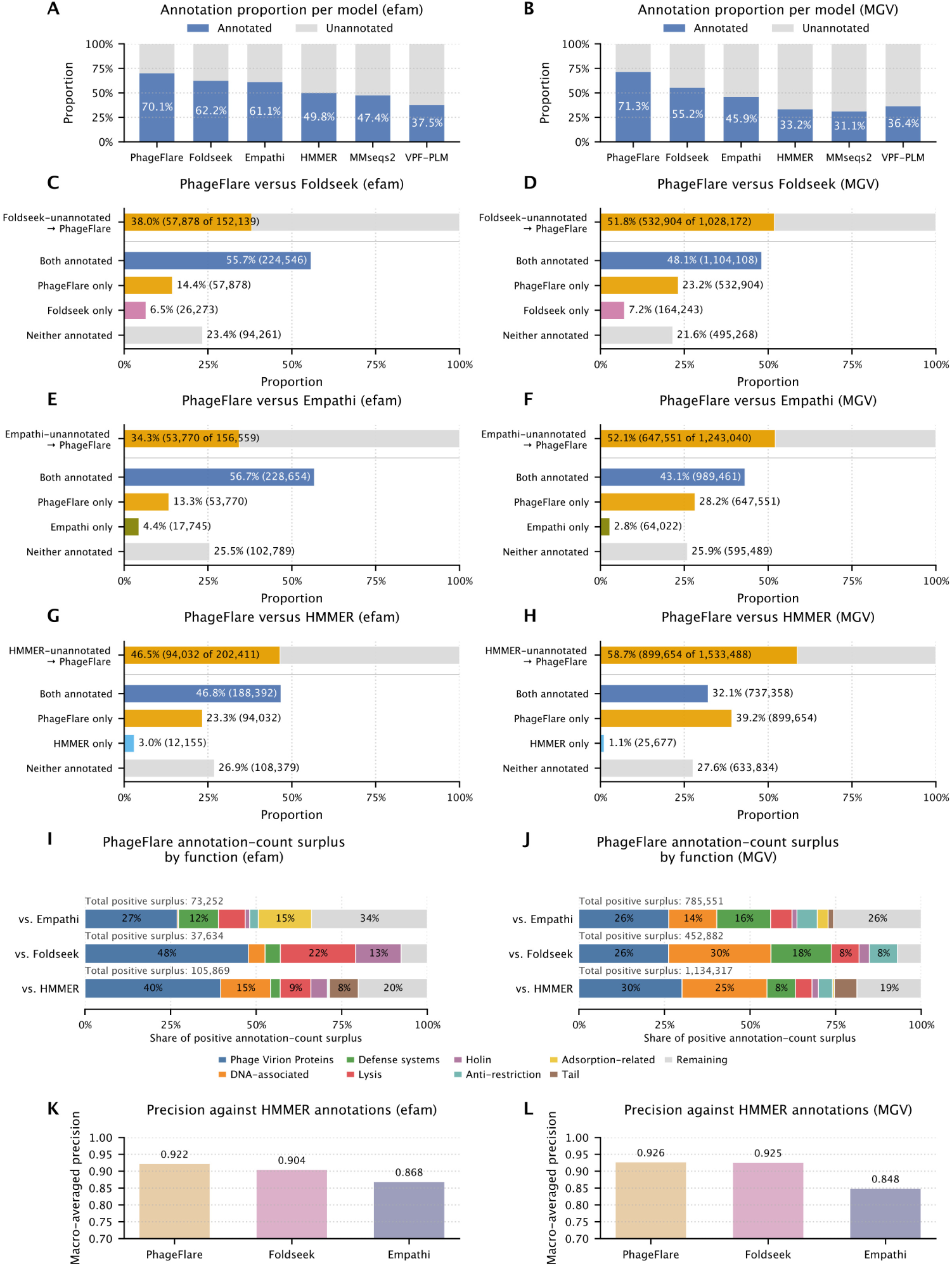
PhageFlare expands functional annotation coverage across the efam and MGV datasets. (**A**, **B**) Proportions of annotated proteins (blue) versus the remaining unannotated (gray) by PhageFlare, Empathi, Foldseek, HMMER, MMseqs2, and VPF-PLM in efam (**A**) and MGV (**B**). Values within the bars indicate the percentage of proteins annotated by each model. (**C**, **D**) Per-protein overlap between annotations assigned by PhageFlare and Foldseek in efam (**C**) and MGV (**D**). The upper bar in each panel represents all proteins unannotated by Foldseek; the orange segment denotes the proportion of these proteins annotated by PhageFlare, whereas the gray segment denotes those remaining unannotated by both models. The four lower bars partition the complete dataset into mutually exclusive groups comprising proteins annotated by both models, only by PhageFlare, only by Foldseek, or by neither model. (**E**, **F**) Per-protein overlap between annotations assigned by PhageFlare and Empathi in efam (**E**) and MGV (**F**). (**G**, **H**) Per-protein overlap between annotations assigned by PhageFlare and HMMER in efam (**G**) and MGV (**H**). (**I**, **J**) Horizontal 100% stacked bars showing the functional composition of the annotation-count surplus produced by PhageFlare relative to Empathi, Foldseek, and HMMER in efam (**I**) and MGV (**J**). For each comparison, positive surplus was calculated as the per-label difference between PhageFlare and comparator annotation counts, retaining only positive differences. The eight labels with the largest mean percentage contributions across all six comparisons are displayed individually in a consistent order and color scheme; all other labels with positive surplus are combined as ’Remaining’. (**K**, **L**) Agreement of PhageFlare, Foldseek and Empathi predictions with HMMER-derived functional annotations in efam (**K**) and MGV (**L**). HMMER annotations were treated as reference labels, and macro-precision values were calculated across 45 functional labels. Values above the bars indicate the corresponding metric estimates.

We further quantified PhageFlare’s capacity to annotate proteins left unannotated by Foldseek (Fig. 5C,D). In efam, 55.7% of the total proteins were annotated by both PhageFlare and Foldseek, whereas 14.4% were annotated only by PhageFlare and 6.5% only by Foldseek. Among the 152,139 proteins unannotated by Foldseek, PhageFlare assigned functional labels to 57,878 (38.0%). This asymmetry was more pronounced in MGV: 48.1% of proteins were annotated by both models, 23.2% only by PhageFlare and 7.2% only by Foldseek. PhageFlare thus annotated 532,904 of the 1,028,172 proteins unresolved by Foldseek (51.8%).

Similar trends were observed relative to Empathi and HMMER. PhageFlare annotated 34.3% of Empathi-unannotated proteins in efam and 52.1% in MGV, whereas the fractions of the complete datasets annotated only by Empathi were 4.4% and 2.8%, respectively (Fig. 5E,F). Among proteins unannotated by HMMER, PhageFlare assigned functional labels to 94,032 of 202,411 proteins in efam (46.5%) and 899,654 of 1,533,488 proteins in MGV (58.7%). By comparison, only 3.0% and 1.1% of the complete efam and MGV datasets, respectively, were annotated exclusively by HMMER (Fig. 5G,H).

The functional composition of PhageFlare’s annotation-count surplus varied across datasets and comparators (Fig. 5I,J). In efam, phage virion proteins contributed the largest share of the positive surplus relative to Empathi (27%), Foldseek (48%) and HMMER (40%). Relative to Foldseek, lysis proteins and holins contributed a further 22% and 13%, respectively. In MGV, DNA-associated proteins and phage virion proteins accounted for 30% and 26% of the positive surplus relative to Foldseek, whereas phage virion proteins and DNA-associated proteins contributed 30% and 25% relative to HMMER. Defense-system proteins also contributed prominently in MGV, accounting for 16% of the positive surplus relative to Empathi and 18% relative to Foldseek. Thus, although several functional labels made substantial contributions, the expansion encompassed multiple structural and non-structural labels.

The expanded annotation coverage was supported by high precision in HMMER-annotated subsets. With HMMER-derived functional annotations as reference labels, PhageFlare maintained higher macro-averaged precision than both Foldseek and Empathi across 45 functional labels for proteins annotated by HMMER. PhageFlare achieved macro-averaged precision values of 0.922 in efam and 0.926 in MGV, compared with 0.904 and 0.925 for Foldseek and 0.868 and 0.848 for Empathi, respectively (Fig. 5K,L). Thus, PhageFlare combined broader annotation coverage with the highest macro-averaged precision among the three models in both datasets, indicating its capacity to extend functional annotation while maintaining stringent prediction criteria.

Sequence length analysis revealed that proteins shorter than 100 amino acids constituted a larger proportion of efam and MGV than of the training dataset (28.2% and 31.4%, respectively, versus 7.6%; Extended Data Fig. 4). This length interval exhibited the lowest annotated-to-unannotated ratio in both external datasets and accounted for 56.3% and 49.8% of proteins remaining unannotated by PhageFlare in efam and MGV, respectively. Thus, despite the broad annotation coverage achieved by PhageFlare, short proteins remained disproportionately represented among unresolved predictions.

Collectively, these analyses connect the evaluation of structure-disjoint proteins with the potential scale of annotation in external phage protein collections. The structure-disjoint benchmark established that PhageFlare retained predictive performance when Foldseek detected no significant structural relationship to the training proteins. The efam and MGV analyses further showed that PhageFlare assigned candidate functions to a substantial proportion of proteins left unresolved by structural alignment and that these annotations spanned diverse functional labels.

### PhageFlare recovers many lysogenic phages misclassified by BACPHLIP

Computational prediction of phage lifestyle often relies on identifying lysogeny-associated genes, including those involved in genome integration and transcriptional repression. Failure to recognize divergent lysogeny-associated proteins can therefore lead to temperate phages being misclassified as lytic [42]. We investigated whether PhageFlare’s enhanced sensitivity in protein function annotation translates into phage lifestyle classification, particularly the recovery of lysogenic phages overlooked by BACPHLIP. For PhageFlare, Empathi, Foldseek and HMMER, a genome was classified as lysogenic when its encoded proteins included at least one protein predicted to perform integration and at least one predicted transcriptional repressor; otherwise, it was classified as lytic. This operational criterion established a uniform, annotation-based benchmarking framework to compare all four models alongside BACPHLIP.

We first evaluated performance on a balanced lifestyle benchmark comprising 6,173 complete inferred lytic phage genomes [52] and 6,173 complete prophage genomes from the MGV dataset. PhageFlare consistently achieved the highest values across the three reported metrics (Fig. 6A). Its *F*_1_ score, MCC and balanced accuracy were 0.790, 0.576 and 0.788, respectively, compared with 0.776, 0.504 and 0.734 for BACPHLIP, 0.777, 0.530 and 0.763 for Empathi, 0.756, 0.462 and 0.722 for Foldseek, and 0.750, 0.447 and 0.713 for HMMER. These results underscore the capacity of PhageFlare in phage lifestyle classification.

**Fig. 6:**
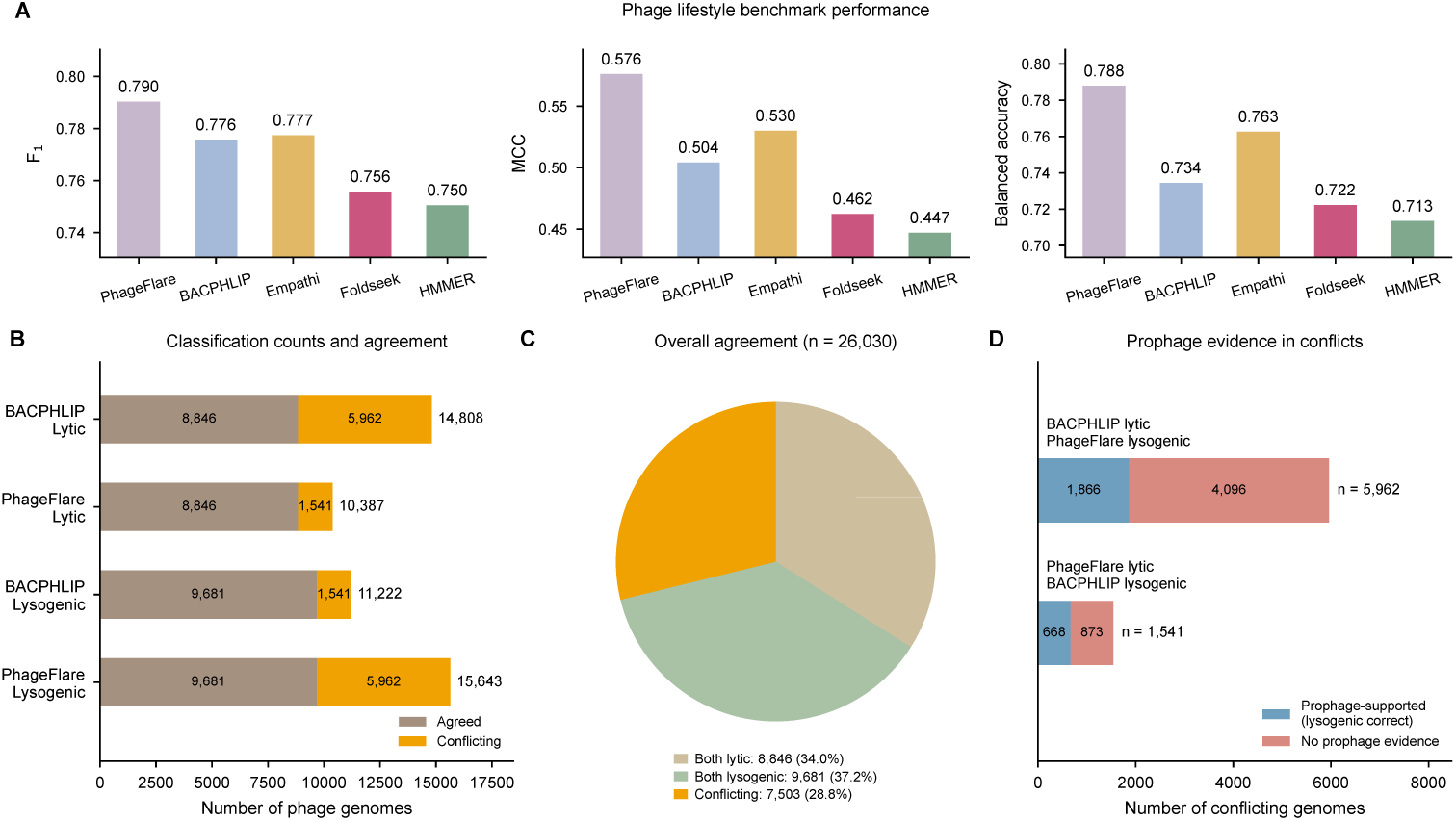
PhageFlare performance in phage lifestyle classification. (**A**) Genome-level classification performance of PhageFlare, BACPHLIP, Empathi, Foldseek and HMMER on the phage lifestyle benchmark, evaluated using the *F*_1_ score, MCC and balanced accuracy. (**B**) Distribution of lytic and lysogenic classifications for complete genomes in MGV by PhageFlare and BACPHLIP. Stacked bars differentiate concordant predictions from conflicting ones. (**C**) Aggregate prediction concordance between PhageFlare and BACPHLIP across complete MGV genomes. (**D**) Analysis of prophage evidence within discordantly classified genomes. Genomes designated as prophages in the MGV metadata are shown in blue and support the lysogenic classification; genomes without a prophage designation are shown in salmon. The absence of a prophage designation was treated as a lack of supporting evidence rather than definitive evidence of a lytic lifestyle.

We next examined lifestyle predictions across 26,030 complete genomes in the MGV, utilizing its pre-computed BACPHLIP predictions. PhageFlare classified 15,643 genomes as lysogenic, compared with 11,222 for BACPHLIP (Fig. 6B). Although the two models exhibited concordance across 18,527 genomes, their predictions differed for 7,503 genomes (28.8%; Fig. 6C). These disagreements were highly asymmetric: PhageFlare assigned a lysogenic lifestyle to 5,962 genomes classified as lytic by BACPHLIP, whereas the reverse disagreement involved 1,541 genomes. Consequently, PhageFlare reassigned 40.3% of BACPHLIP’s 14,808 lytic predictions to the lysogenic category, identifying a substantial set of potentially overlooked lysogenic phages.

To distinguish supported recoveries from unresolved prediction disagreements, we examined prophage designations in the MGV metadata (Fig. 6D). Among the 5,962 genomes classified as lytic by BACPHLIP but lysogenic by PhageFlare, 1,866 were designated as prophages, providing supporting evidence for the PhageFlare predictions. These recoveries represented 12.6% of all BACPHLIP lytic predictions in the complete MGV collection. Conversely, 668 prophage-designated genomes were classified as lysogenic by BACPHLIP but lytic by PhageFlare. PhageFlare therefore recovered approximately 2.8 times as many prophage-supported lysogenic genomes missed by BACPHLIP as were recovered in the reverse comparison. The remaining 4,096 and 873 genomes in the respective disagreement groups lacked prophage designations. However, their lifestyles remained unresolved because the absence of a prophage annotation does not establish a lytic lifestyle.

Recovery of these overlooked lysogenic genomes suggests a potential application of PhageFlare in screening phages for therapy, where obligately lytic candidates are generally preferred and lysogeny-associated genes are assessed during genomic characterization [53]. PhageFlare could provide a complementary screening step by flagging apparently lytic candidates whose predicted protein functions indicate lysogenic potential. The MGV results support this use for candidate assessment, although its performance requires evaluation in collections of therapeutic isolates. Furthermore, we note that a lytic prediction under the present operational criterion does not independently guarantee obligately lytic growth or therapeutic suitability. Experimental lifestyle characterization and broader genomic assessment remain necessary [54].

## Discussion

The expanding catalog of phage sequences has exposed a broadening gap between viral discovery and functional annotation. Although structural alignment has improved annotation sensitivity, a large proportion of proteins remain functionally uncharacterized because they lack significant structural matches to known references [29]. Therefore, we developed PhageFlare, a deep learning framework specifically designed to illuminate this functional dark matter beyond the reach of structural alignments. To explicitly evaluate its capability, we established three structure-disjoint benchmarks across varying *E*-value thresholds (10*^−^*^2^, 10*^−^*^3^, 10*^−^*^4^), where PhageFlare achieved macro-averaged *F*_1_ scores of 0.738, 0.871, and 0.889, respectively. This demonstrates PhageFlare’s capacity to predict functions for phage proteins unresolved by the evaluated structural-alignment-based approach. We further evaluated the influence of the PLM backbone on predictive performance, finding that backbone selection has substantial impact. Among the three evaluated PLM backbones (ESM-C, ESM-2, ProtBert), PhageFlare (ESM-C) achieved the best performance, followed by PhageFlare (ESM-2), whereas PhageFlare (ProtBert) exhibited a large performance gap. In addition, we included other PLM-based models in the benchmark and confirmed that PhageFlare outperformed the strongest competitor, Empathi (tuned thresholds), which attained macro-averaged *F*_1_ scores of 0.732, 0.789, and 0.814, respectively.

To investigate patterns of PhageFlare latent embeddings, we analyzed local structural features of the major capsid and portal proteins. The observed co-localization of divergent training and test proteins within classifier projections coincided with the preservation of specific local helical elements, despite limited overall structural similarity. These observations indicate that proximity in latent embedding space can reflect shared local structural elements, even in the absence of overall structural similarity.

To evaluate whether PhageFlare’s benchmark performance translates to large-scale discovery, we applied it to the efam and MGV datasets. In these collections, PhageFlare assigned candidate functions to 38.0% and 51.8% of proteins that remained unannotated by Foldseek, respectively. Furthermore, these new functional annotations spanned diverse biological processes, including virion morphogenesis, nucleic acid metabolism, host defense, transcriptional regulation, and lysis, providing a framework to prioritize targets for downstream investigation. Although the sparsity of experimentally validated functional annotations limits direct assessment of predictive performance in these datasets, PhageFlare’s expanded coverage was accompanied by high reference-based precision. Using HMMER-derived functional annotations as reference labels, PhageFlare achieved macro-averaged precision values of 0.922 in efam and 0.926 in MGV across 45 functional labels within the HMMER-annotated subsets, compared with 0.904 and 0.925 for Foldseek and 0.868 and 0.848 for Empathi, respectively (Fig. 5K,L).

Beyond assigning protein functions, we investigated the utility of PhageFlare in phage lifestyle prediction. Using predicted integration and transcriptional repressor functions, PhageFlare achieved better classification performance than the evaluated alternatives. In MGV, it identified lysogenic potential in 1,866 prophage-designated genomes that were misclassified as lytic by BACPHLIP, compared with 668 recovered in the reverse comparison (Fig. 6). This could provide a complementary screening step for prioritizing therapeutic phage candidates by flagging apparently lytic genomes that encode lysogeny-associated functions.

Several limitations bound the current scope of PhageFlare. First, the framework operates on a fixed ontology of 45 functional labels, restricting its predictive capacity to the granularity and scope of existing annotations. Second, the precision values of PhageFlare on HMMER-annotated subsets of efam and MGV do not establish its precision across the complete datasets. Nevertheless, in the absence of ground-truth functional annotations, these subsets provide a practical basis for estimating precision. Third, PhageFlare is not designed as a standalone diagnostic tool for phage lifestyle classification. Specifically, the predicted presence of integration and transcriptional repressor proteins does not definitively guarantee a lysogenic lifestyle. Therefore, PhageFlare is best used as a complementary screening step that flags potentially lysogenic candidates within pools of putatively lytic phages.

A specific priority for future development is the annotation of short proteins (fewer than 100 amino acids). These short sequences are disproportionately represented in different environmental datasets and constitute approximately half of the proteins that PhageFlare left unannotated. Although some of these unresolved sequences may represent false-positive gene predictions, given that short open reading frames frequently occur by chance and are difficult to differentiate from genuine protein-coding genes [55], a previous study uncovered a vast repertoire of functionally relevant small proteins [56]. Consequently, their disproportionate representation among unresolved predictions underscores the critical need to expand collections of curated short proteins, which will be essential for extending annotation coverage into this challenging sequence space.

In conclusion, PhageFlare extends the annotation of phage functional dark matter to proteins that remain unresolved by structural-alignment-based retrieval against the same reference set. It provides complementary functional evidence where significant structural matches to annotated references are unavailable. Its applications to metagenomic datasets demonstrate how this capability can broaden functional annotation coverage across diverse phage proteins. PhageFlare thus represents a step towards next-generation phage annotation tools that extend functional inference beyond structural-alignment-based approaches, enabling further exploration of previously unresolved phage functional space.

## Methods

### Sequence-disjoint dataset preparation

Protein sequences were collected from PHROGs v4 [57] and the 14 April 2025 release of INPHARED [58]. PHROGs entries assigned to the ’unknown function’ category were excluded. Across both resources, unannotated proteins and proteins longer than 2,048 amino acids, or containing the noncanonical symbols ’X’, ’B’, ’Z’, ’J’, or the stop-codon marker ’*’ were removed from the dataset. Functional annotations from both resources were harmonized to 45 functional labels according to the Empathi mapping scheme [33]. Exact duplicate protein sequences were collapsed within and across the two resources, while retaining the union of their functional labels.

The resulting proteins were clustered using MMseqs2 [59], with a minimum sequence identity of 0.3, a minimum coverage of 0.8, and an E-value threshold of 10*^−^*^3^. To account for functional annotation heterogeneity among proteins within the same cluster, original protein-level labels were preserved for evaluation. During model training, annotations shared universally across all members of a cluster were defined as positive targets; conversely, labels present in only a subset of cluster members were designated as inconsistent and masked from the functional classification loss calculation. ’Sequence-disjoint’ denotes that each MMseqs2 cluster was assigned exclusively to the training, validation or test dataset. Consequently, no proteins in the validation and test datasets shared a cluster with a training protein under the specified identity, coverage, and E-value criteria.

Clusters were divided among the training, validation, and test datasets in an 80:10:10 ratio. The final dataset contained 377,115 training, 66,358 validation, and 58,519 test proteins. Genomic context was available only for INPHARED-derived samples, whereas PHROGs-derived samples comprised target proteins only. For each INPHARED target protein, up to six genomic-context proteins, corresponding to three upstream and three downstream genes, were included, with unavailable genomic-context proteins represented by empty entries.

### Structure-disjoint benchmark preparation

The structure-disjoint benchmark was prepared from the same harmonized PHROGs– INPHARED protein collection and MMseqs2 clusters. A fixed held-out pool of 1,200 clusters was randomly selected, and all remaining clusters were initially assigned to the training pool. This procedure yielded 32,662 held-out proteins and 469,330 candidate training proteins.

Each protein was represented for structural searching using a PDB-format structure from the Phold structural database, when available. For proteins without an available structure, ProstT5-derived structural representations were generated by Foldseek. Separate Foldseek databases were constructed for the PDB- and ProstT5-represented subsets. Four database-level comparisons spanning the Cartesian product of the query and target subsets were performed: PDB-query–PDB-target, PDB-query–ProstT5-target, ProstT5-query–PDB-target, and ProstT5-query–ProstT5-target. Thus, each protein occurred in only one query database and was searched against the two target databases. We used exhaustive searches in Foldseek alignment with alignment type 2 and three different E-value thresholds of 10*^−^*^2^, 10*^−^*^3^, and 10*^−^*^4^, respectively. Any training protein returned as a significant hit for a held-out query in any of the four searches was removed.

The training databases were rebuilt after each filtering step, and the search-and-removal procedure was repeated until a verification search returned no significant held-out-to-training alignments. Four successive filtering rounds reduced the size of the candidate training set. Because the query–target direction may affect Foldseek alignment results, reverse-direction searches were subsequently performed using the training proteins as queries and the fixed held-out proteins as targets. An explicit repeat search of the resulting dataset detected no significant alignment in any representation combination. Following this procedure, three distinct structure-disjoint benchmarks were generated corresponding to *E*-value thresholds of 10*^−^*^2^, 10*^−^*^3^, and 10*^−^*^4^, respectively.

’Structure-disjoint’ denotes that no protein in the training dataset produced a significant Foldseek alignment to any protein in the held-out pool, or vice versa, across the four PDB/ProstT5 representation combinations at the specified E-value threshold. This definition applies to the separation of the training dataset from the complete held-out pool. The held-out pool was randomly partitioned at the protein level into equally sized validation and test datasets. Because structural filtering was performed against the complete held-out pool before this division, both validation and test datasets remained structure-disjoint from the final training dataset. Up to six genomic-context proteins were added to INPHARED-derived samples as described above.

### Efam and MGV data collection

For efam, we used the dereplicated and filtered efam-XC protein set. This dataset comprised target proteins only and did not include genomic context. The MGV dataset was obtained from release v1.0, dated 8 July 2021. Genomes whose ’checkv quality’ metadata field was exactly ’Complete’ were retained, resulting in 26,030 genomes. Rather than repredicting genes, we retrieved the corresponding proteins from the protein FASTA file distributed with MGV. To incorporate genomic context, each MGV protein was paired with up to six genomic-context proteins corresponding to three upstream genes and three downstream genes.

### Phage lifestyle benchmark preparation

A phage lifestyle benchmark was constructed from inferred lytic phages reported in a previous study [52] and complete prophages from MGV. For lytic phage genomes, protein-coding genes were predicted using prodigal-gv [12, 60] in metage-nomic mode. For the lysogenic class, MGV genomes were retained only when both the ’checkv quality’ field was ’Complete’ and the ’prophage’ field was ’Yes’. This filtering produced 6,173 prophage genomes and 512,400 release-provided protein sequences.

Because the lytic phage collection was substantially larger, it was downsampled to 6,173 genomes. The selected lytic phage genomes were combined with 6,173 MGV prophage genomes, producing a balanced dataset of 12,346 genomes. Genome-level labels were encoded as 1 for lytic phages and 0 for lysogenic phages. Up to three proteins on either side of each target protein were included as genomic-context proteins. The resulting dataset was used for the phage lifestyle benchmark.

### PhageFlare architecture

PhageFlare is a multilabel classifier that models function-specific patterns of residue contribution and uses the predicted functions of proteins in the genomic neighborhood to refine the functional annotation of a target protein. For a target protein and *M* genomic-context proteins, a shared-weight predictor was applied independently to each protein sequence. The implementation accepted up to six genomic-context proteins (*M ≤* 6), corresponding to the three upstream and three downstream genes. Missing genomic-context proteins were represented by empty inputs and excluded using a validity mask.

Each sequence was tokenized with the ESM sequence tokenizer and processed by the frozen 600-million-parameter ESM-C encoder. ESM-C produced a stack of latent embeddings, *E ∈* R^36^*^×T^ ^×H^* , where *T* denotes sequence length and *H* = 1,152 is the ESM-C hidden dimension. Two independent learned linear projections across the 36-layer axis generated two residue embeddings for domain prediction and protein function prediction, respectively. A BiLSTM converted the residue embeddings (domain) into residue-level domain probabilities. The residue embeddings (function) and domain probabilities were subsequently processed by the function-conditioned residue pooling module (Supplementary Notes) to generate a unified embedding **u***_c_ ∈* R*^H^* for each function *c*. After layer normalization, each **u***_c_* was passed through a separate two-layer MLP, comprising a GELU activation, to produce the logit *s_c_*. The resulting logits were assembled into the function logit vector **s** = (*s*_1_*, . . . , s_C_*) *∈* R*^C^*. Thus, all functions shared the ESM-C encoder and attention projections, while retaining function-specific queries and MLP classifiers.

The shared predictor produced a raw target function logit vector **s**_0_ *∈* R*^C^* for the target protein and a corresponding logit vector **s***_m_ ∈* R*^C^* for each valid genomic-context protein *m*. These logits, together with their corresponding relative gene offset, were passed to the genomic-context module (Supplementary Notes). This module generated context correction logits and a per-function gate, which together defined a gated residual correction to the raw target function logits. An elementwise sigmoid was then applied to the corrected logits to obtain the final target function probabilities. When no valid genomic context was available, the context correction was omitted and PhageFlare used the raw target function logits.

### PhageFlare training

PhageFlare was trained using AdamW [61] with a peak learning rate of 5 *×* 10*^−^*^4^, *β*_1_ = 0.9, *β*_2_ = 0.95 and weight decay 0.01. The learning rate increased linearly during the first 10% of optimization steps and then decayed linearly to 1% of its peak value. The ESM-C encoder remained frozen throughout training. Gradients were clipped to a maximum *ℓ*_2_ norm of 1.0.

For protein function classification, the model was optimized using an asymmetric multilabel loss applied independently to the *C* functional-label logits. For protein *i* and functional label *c*, *z_ic_* denotes the predicted logit and *y_ic_* denotes its target, where *y_ic_* = 1 indicates that the function is present, and *y_ic_* = 0 indicates that it is absent. The set of function-supervised protein–label pairs is denoted as

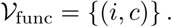

The function-classification loss was

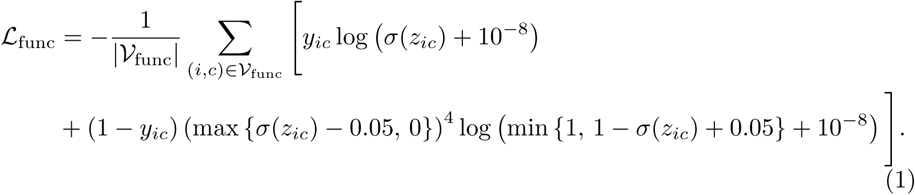

Here, *σ*(*·*) denotes the sigmoid function. The first term is the positive-label binary cross-entropy because the positive focusing parameter is *γ*_+_ = 0. In the second term, the negative-label probability 1 *− σ*(*z_ic_*) is increased by 0.05 and capped at 1, while the corresponding negative focusing factor is raised to *γ_−_* = 4.

For residue-level domain classification, the training dataset comprised domain-annotated bacterial and non-bacteriophage viral proteins from UniProt [62]. Let *I*_dom_ denote the set of proteins with domain annotations and let *R_i_* denote the set of valid amino acid residues in protein *i*. *bos*, *eos*, and padding positions were excluded from *R_i_*. For residue *r ∈ R_i_*, the target *d_ir_ ∈ {*0, 1*}* indicated whether the residue was inside a domain, and *d̂_ir_* was the predicted in-domain probability. For numerical stability, the predicted probability was clipped as

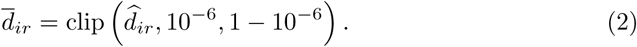

For each domain-annotated protein, a per-protein scaling factor was calculated from the relative numbers of residues outside and inside annotated domains:

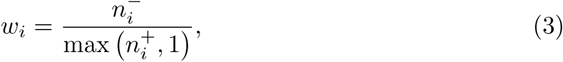

where *n*^+^_*i*_ and *n*^−^_*i*_ denote the numbers of valid amino-acid residues inside and outside annotated domains, respectively. The domain-classification loss was obtained by averaging binary cross-entropy over the valid residues of each annotated protein, multiplying the resulting per-protein mean by *w_i_*, and then averaging over domain-annotated proteins:

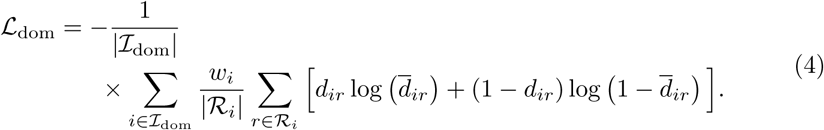

All function-classification samples were excluded from the domain-classification loss.

The function- and domain-classification objectives were combined while limiting the contribution of the auxiliary domain loss to at most one-third of the contemporaneous function-classification loss:

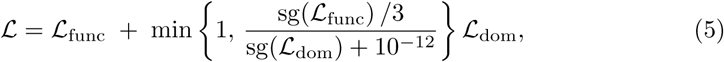

where sg(*·*) denotes the stop-gradient operation, which was applied only when computing the ratio.

A checkpoint was saved after each training epoch. The best epoch was selected as that with the highest mean label-wise validation AUPRC, calculated from the prediction probabilities. For each epoch, an initial binarization threshold was estimated independently for every functional label by evaluating the exact breakpoints defined by its unique validation-set probabilities and selecting the threshold that maximized its label-specific *F*_1_ score. Independently binarizing hierarchically related labels could, however, produce inconsistent predictions in which a child label was positive while its parent was negative. Hierarchical consistency was therefore enforced recursively by suppressing a positive child prediction whenever its parent was negative, such that a positive descendant was retained only when all of its ancestors were also positive. For each functional label with a parent or child label, candidate thresholds were drawn from the exact validation-probability breakpoints, including a value immediately above the maximum probability, while the thresholds for all other labels were held fixed. An update was accepted only when it strictly increased the hierarchy-adjusted macro-*F*_1_ score. Optimization stopped when a complete pass produced no improvement or after ten passes. The checkpoint from the selected epoch and the hierarchy-aware thresholds estimated for that epoch were subsequently used for test set evaluation, with a label called positive when its predicted probability was greater than or equal to its threshold.

For the structure-disjoint benchmarks, PhageFlare was trained on the corresponding benchmark-specific training datasets. For inference on the efam, MGV, and phage lifestyle benchmark, PhageFlare used the model checkpoint trained on the sequence-disjoint training dataset.

### Training and testing competing models

For the structure-disjoint benchmarks, Empathi was trained on the corresponding benchmark-specific training datasets across 45 functional labels before evaluation. Two versions of Empathi were evaluated: Empathi, for which a universal binarization threshold of 0.5 was applied to all functional labels, and Empathi (tuned thresholds), for which function-specific thresholds were optimized on the validation dataset using the *F*_1_ score. For HMMER and DHR, the reference database comprised the training dataset of the corresponding benchmark, and the query database comprised its test dataset. VPF-PLM and GOPhage were excluded from the benchmarks as they have different output functional labels.

For inference on the efam and MGV datasets, Empathi and VPF-PLM were used with their respective model checkpoints. The binarization threshold for Empathi was set to its recommended stringent value of 0.95, whereas the function-specific binarization thresholds for VPF-PLM were those calibrated for efam and provided with its accompanying script. For Foldseek, HMMER, and MMseqs2, the reference database comprised the sequence-disjoint training dataset. The *E*-value thresholds of Fold-seek were 0.005 and 0.05 for efam and MGV, respectively. The *E*-value thresholds of HMMER and MMseqs2 were 10*^−^*^3^ for both datasets. GOPhage was excluded from these experiments due to the prohibitive storage overhead generated by its large intermediate files when processing large-scale datasets.

For the phage lifestyle benchmark, Empathi was used with its own model check-point and its recommended stringent binarization threshold of 0.95. The reference database for Foldseek and HMMER comprised the sequence-disjoint training dataset. BACPHLIP was used with a binarization threshold of 0.5. The *E*-value thresholds of Foldseek and HMMER were 10*^−^*^3^.

For Foldseek and MMseqs2, a query protein was assigned the functional labels of the protein in the reference database with the lowest *E*-value, contingent upon the corresponding *E*-value threshold. Queries lacking reference hits below this significance threshold were assigned an ’unknown’ label. For HMMER, query proteins were searched against the reference database using phmmer, and all functional labels of the reference protein with the lowest full-sequence *E*-value were transferred to the query, provided that the *E*-value was less than or equal to the threshold. Queries without an accepted reference hit were assigned an ’unknown’ label. For DHR, functional labels were assigned based on the nearest reference protein in the embedding space. Because DHR unconditionally assigns labels to all query proteins, it was excluded from the functional annotation coverage experiments on the efam and MGV datasets. The specific version used for each method is detailed in Supplementary Table 1. Other hyper-parameter settings of HMMER, MMseqs2, Foldseek are listed in Supplementary Table 2.

### Benchmark evaluation

The function prediction benchmark comprised multilabel classification tasks. We evaluated the predictive performance of each model using *F*_1_ score, AUPRC and MCC. AUPRC was computed using *scikit-learn* [63]. For HMMER and DHR, their AUPRC values were calculated based on binary prediction labels. *F*_1_ score and MCC were calculated as

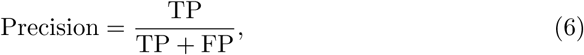

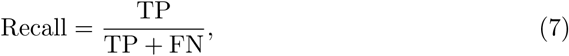

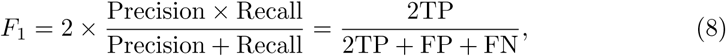

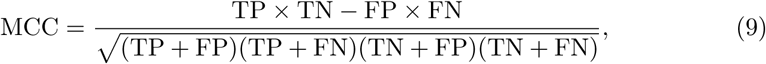

where TP, TN, FP, and FN denote the numbers of true positives, true negatives, false positives, and false negatives, respectively.

The phage lifestyle benchmark comprised a binary classification task. Given the balanced class distribution, model performance was evaluated using the *F*_1_ score, MCC, and balanced accuracy.

### Train-test co-localization analysis in the PhageFlare latent space

To assess whether structure-disjoint training and test proteins of a functional label were similar in the PhageFlare latent space, we analyzed each functional label independently using the PhageFlare model trained on the structure-disjoint benchmark. For each functional label, we extracted the 576-dimensional embedding of the target protein post-GELU activation, following the first linear layer and before the second linear layer, in the corresponding per-function MLP classifier. Each analysis included all positive training and test samples for the corresponding function, together with up to 2,000 negative samples that were sampled from the pooled training and test sets.

Dimensionality reduction was performed independently for each function. The 576-dimensional embeddings were first reduced to 50 principal components using randomized singular-value decomposition and were then projected into two dimensions using t-SNE. Separate projections were generated with a perplexity of 30. t-SNE was run with principal component analysis (PCA) initialization, an automatically determined learning rate, an early-exaggeration factor of 12, the Barnes–Hut approximation and 2,000 optimization iterations. The samples from training and test datasets were subsequently distinguished in the resulting two-dimensional plots to evaluate their spatial co-localization.

The structures of the analyzed proteins were predicted as single chains using AlphaFold 3 [64]. Pairwise structural similarity was quantified for all train–test combinations using US-align [47]. For each comparison, TM-scores were normalized independently by the lengths of the first and second proteins, and their arithmetic mean was reported as ’TM-score mean’. RMSD was calculated over the structurally aligned residues. Coverage was defined as the aligned length divided by the full length of the corresponding protein, and aligned sequence identity was defined as the proportion of identical residue pairs within the US-align alignment. Protein structures and structural superpositions were visualized using PyMOL.

### PhageFlare latent embedding analysis for class separability

To assess the separability of functional labels in the latent space learned by PhageFlare, we extracted embeddings for all proteins in the test dataset of the structure-disjoint benchmark with the *E*-value threshold of 10*^−^*^3^, using the PhageFlare model trained on the training dataset of this benchmark. For each functional label, we retrieved the 576-dimensional embedding post-GELU activation, following the first linear layer, in the corresponding per-function MLP classifier. The t-SNE projection settings were identical to the protocol described above. The resulting projections were visualized by coloring proteins according to their binary ground-truth labels.

## Data availability

The PHROGs v4 database is available at https://phrogs.lmge.uca.fr/, INPHARED (14 April 2025 release) is available at https://github.com/RyanCook94/inphared, efam is available at https://dc.cyverse.org/dataset/efam, and MGV is available at https://portal.nersc.gov/MGV/. The 6,173 lytic phage genomes are available at https://doi.org/10.6084/m9.figshare.28911734. Domain annotations are available at https://www.uniprot.org/.

## Code availability

PhageFlare source code is available at https://github.com/ericcombiolab/PhageFlare. ESM-C is available at https://github.com/Biohub/esm. ESM-2 is available at https://github.com/facebookresearch/ESM. ProtBert is available at https://huggingface.co/Rostlab/protbert.

## Acknowledgements

Fig. 1A was created with BioRender.com. The project is supported by the Young Collaborative Research grant (No. C2004-23Y, No. C2005-24Y), HMRF grant (No. 11221026), and HKBU RCMS (RCMS/24-25/03).

## Author contributions

C.W., L.Z. (Louxin) and L.Z. (Lu) conceived the project. C.W. designed the model and performed the experiments. C.W., Y.X., Z.L., L.Z. (Louxin), and L.Z. (Lu) contributed to discussions. C.W. wrote the paper. C.W., Y.X., Z.L., L.Z. (Louxin), and L.Z. (Lu) reviewed the paper. L.Z. (Louxin) and L.Z. (Lu) supervised the project.

---

**Extended Data Fig. 1:**
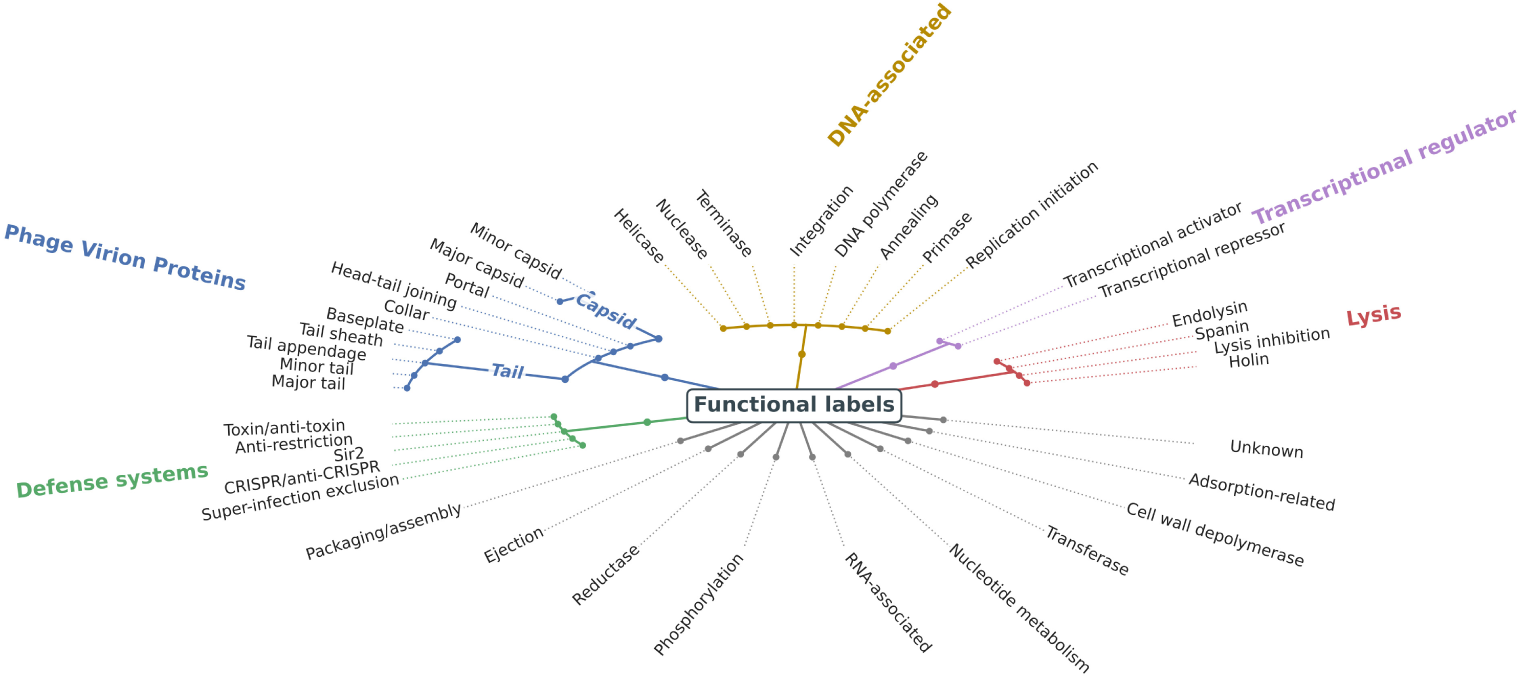
PhageFlare employs 45 hierarchical functional labels derived from Empathi. Proteins for which all functional labels yield negative predictions are designated as ’Unknown’.

**Extended Data Fig. 2:**
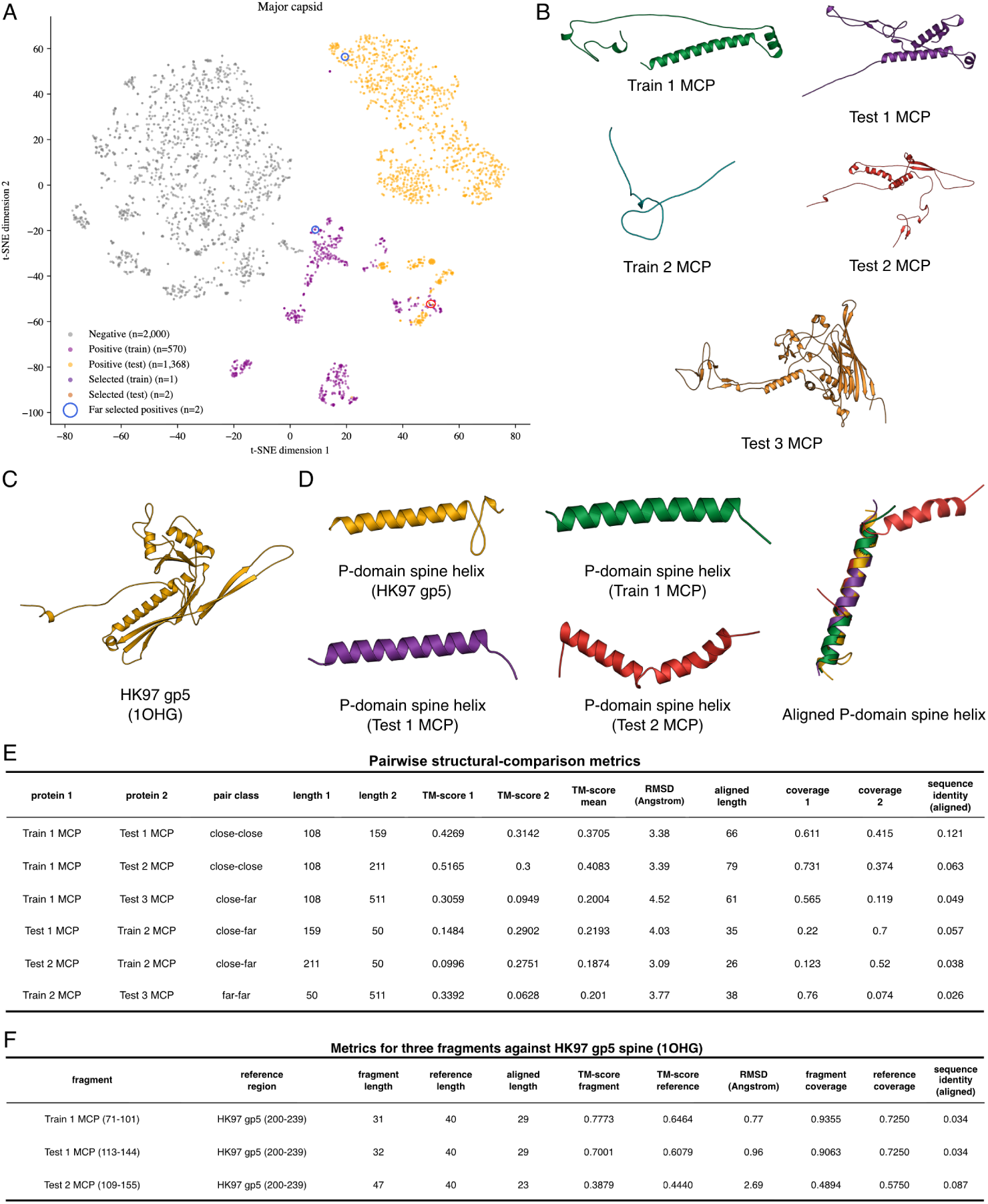
Embedding-guided analysis of major capsid proteins in the structure-disjoint benchmark. (**A**) Two-dimensional t-SNE projection of embeddings from PhageFlare major capsid protein classifier, comprising 570 training positives, 1,368 test positives and 2,000 sampled proteins lacking the major capsid label. One training sample and two test samples positioned close together (Train 1, Test 1 and Test 2) and two more distant samples (Train 2 and Test 3) were selected for structural analysis. (**B**) Single-chain structures of the five selected samples predicted by AlphaFold 3. (**C**) A subunit from the experimentally determined HK97 capsid used as a structural reference (PDB 1OHG). (**D**) P-domain spine helix region from the three embedding-proximal proteins and the reference spine helix. The final subpanel shows their US-align-derived superpositions onto the reference. (**E**) Metrics for all six comparisons between the two training and three test samples. Pair classes describe their selection from proximal or distant positions in the t-SNE projection. (**F**) Metrics for comparisons of the three representative helical regions with the HK97 reference spine.

**Extended Data Fig. 3:**
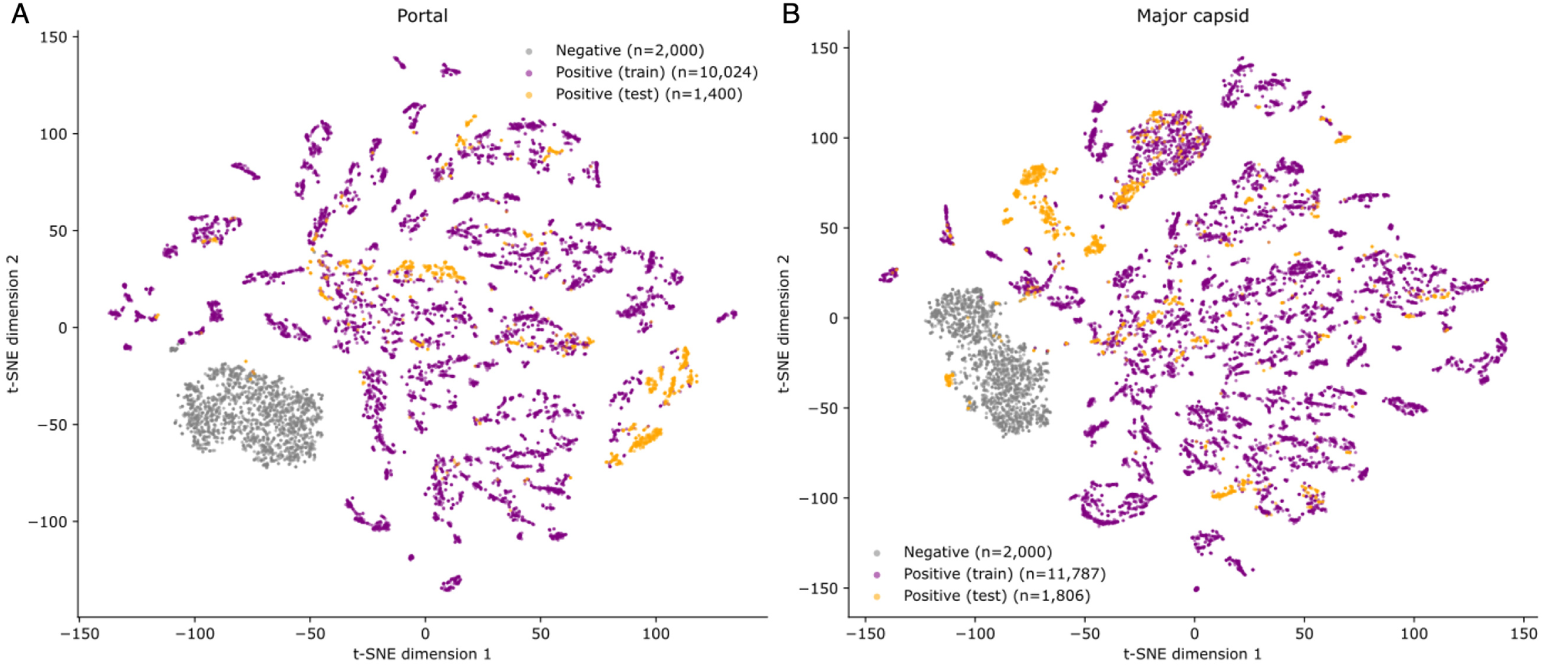
Two-dimensional t-SNE visualization of PhageFlare embeddings in the sequence-disjoint dataset. Projections illustrate protein embeddings generated by the PhageFlare portal classifier (**A**) and the PhageFlare major capsid protein classifier (**B**). Embeddings were obtained from the corresponding per-function MLP classifiers of the model trained on the sequence-disjoint training dataset. Each point represents one protein. Training positives, test positives and sampled proteins lacking the corresponding functional label are distinguished by color.

**Extended Data Fig. 4:**
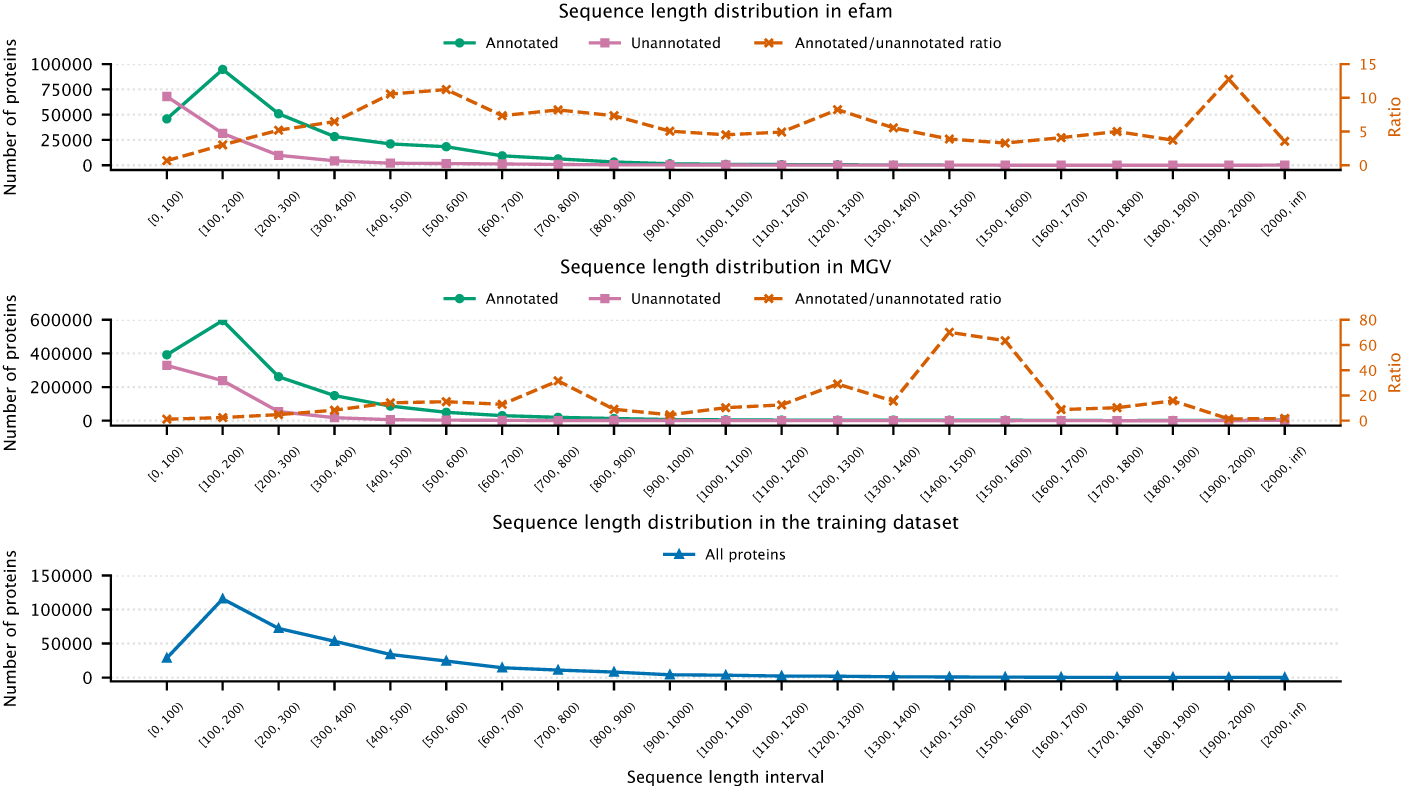
Sequence length distributions in efam, MGV, and the PhageFlare training dataset. Upper and middle panels show the numbers of proteins annotated (green circles) and unannotated (pink squares) by PhageFlare in efam and MGV, respectively. Orange dashed lines indicate the ratio of annotated-to-unannotated proteins within each length interval, plotted against the right-hand axis. The lower panel shows the sequence length distribution of all proteins in the sequence-disjoint training dataset.

