## Supplementary Notes for "Illuminating phage dark matter beyond the reach of structural alignments with PhageFlare"

#### Contents

**1 Supplementary Notes** **1**
1.1 Function-conditioned residue pooling module . . . . . . . . 1
1.2 Genomic-context module . . . . . . . . 2

**2 Supplementary Tables** **3**

**3 Supplementary Figures** **4**

### 1 Supplementary Notes

#### 1.1 Function-conditioned residue pooling module

PhageFlare used distinct learned mixtures of ESM-C embeddings from all layers for the auxiliary domain prediction task and the main function prediction task. Let $\mathbf{E}_\ell \in \mathbb{R}^{T\times H}$ denote the residue-representation matrix from ESM-C layer $\ell$. The domain and function embeddings were

$$\mathbf{D} = \sum_{\ell=1}^{36} a_\ell^{\mathrm{dom}} \mathbf{E}_\ell, \qquad \mathbf{F} = \sum_{\ell=1}^{36} a_\ell^{\mathrm{fun}} \mathbf{E}_\ell, \tag{1}$$

where $a_\ell^{\mathrm{dom}}$ and $a_\ell^{\mathrm{fun}}$ are independently learned scalar coefficients. A one-layer BiLSTM with hidden size $H/2$ in each direction mapped $\mathbf{D}$ to a domain probability $\rho_t \in [0,1]$ for each residue $t$. These probabilities served both as the output of the auxiliary domain-prediction head and as an attention prior for the multi-head attention mechanism in the function-conditioned residue pooling module.

To derive function-conditioned embeddings, PhageFlare learned one query $\mathbf{q}_c \in \mathbb{R}^H$ for each function $c \in \{1,\ldots,C\}$. The queries attended to $\mathbf{F}$ using eight-head scaled dot-product attention. For head $h$, the domain prior was incorporated by adding its logarithm to the query–key compatibility score:

$$A_{ct}^{(h)} = \operatorname*{softmax}_t \left[ \frac{\left(\mathbf{q}_c^{(h)}\right)^\mathsf{T} \mathbf{k}_t^{(h)}}{\sqrt{H/8}} + \log\left(max\left(\rho_t, \epsilon\right)\right) \right], \tag{2}$$

where the *bos*, *eos* and padding-token positions were masked and $\epsilon = 10^{-6}$ ensured numerical stability.

For each attention head, the function-conditioned embedding was computed as:

$$\mathbf{z}_c^{(h)} = \sum_{t\in\mathcal{R}} A_{ct}^{(h)} \mathbf{v}_t^{(h)}, \tag{3}$$

where $\mathbf{v}_t^{(h)}$ denotes the value representation of residue $t$ from residue embeddings (function) and $\mathcal{R}$ is the set of valid residue positions. The outputs of the eight heads were concatenated and linearly projected to obtain the final function-conditioned embedding:

$$\mathbf{z}_c = \mathbf{W}^O \, \text{Concat}\left(\mathbf{z}_c^{(1)}, \ldots, \mathbf{z}_c^{(8)}\right), \qquad \mathbf{z}_c \in \mathbb{R}^H, \tag{4}$$

where $\mathbf{W}^O$ is the output projection of the multi-head attention module.

In parallel, masked mean pooling of $\mathbf{F}$ over valid residue positions, likewise excluding *bos*, *eos* and padding tokens, produced a global embedding $\bar{\mathbf{f}} \in \mathbb{R}^H$. A learned feature-wise gating parameter $\boldsymbol{\gamma} \in \mathbb{R}^H$ combined the two representations:

$$\mathbf{u}_c = \bar{\mathbf{f}} + \sigma(\boldsymbol{\gamma}) \odot \mathbf{z}_c, \tag{5}$$

where $\mathbf{u}_c \in \mathbb{R}^H$ is the unified embedding for function $c$, $\sigma(\cdot)$ denotes the elementwise sigmoid function and $\odot$ denotes elementwise multiplication.

#### 1.2 Genomic-context module

The genomic context module refined the target prediction by embedding the raw target function probabilities and $M$ genomic-context function probabilities into a 128-dimensional latent space, denoted $H_{\text{ctx}} = 128$. Specifically, for protein $m$, the probability vector $\mathbf{p}_m = \sigma(\mathbf{s}_m)$ was projected with a learned matrix $\mathbf{E}_{\text{in}} \in \mathbb{R}^{C \times H_{\text{ctx}}}$. Each genomic-context protein representation was combined with a learned embedding of its relative position $r_m \in \{-3, -2, -1, +1, +2, +3\}$, where negative and positive values denote upstream and downstream positions, respectively. The target was represented analogously and inserted at position $r_0 = 0$. A single-layer BiLSTM then encoded the ordered target and genomic-context latent embeddings in both genomic directions.

Before contextual encoding, a relevance prior $\alpha_m$ was computed for each genomic-context protein $m$. Let $\mathbf{p}_m = \sigma(\mathbf{s}_m) \in [0,1]^C$ and $\mathbf{p}_0 = \sigma(\mathbf{s}_0) \in [0,1]^C$ denote the function probabilities of genomic-context protein $m$ and the target protein, respectively. Their representations in the 128-dimensional latent space were

$$\mathbf{h}_m = \text{LayerNorm}\left(\mathbf{E}_{\text{in}}^{\top} \mathbf{p}_m\right), \qquad \mathbf{h}_0 = \text{LayerNorm}\left(\mathbf{E}_{\text{in}}^{\top} \mathbf{p}_0\right). \tag{6}$$

The uncertainty of the prediction for genomic-context protein $m$ was summarized by its mean Bernoulli entropy,

$$\bar{p}_{mc} = \text{clip}(p_{mc}, \epsilon, 1 - \epsilon), \tag{7}$$

$$\mathcal{H}_m = -\frac{1}{C} \sum_{c=1}^{C} \left[\bar{p}_{mc} \log \bar{p}_{mc} + (1 - \bar{p}_{mc}) \log (1 - \bar{p}_{mc})\right]. \tag{8}$$

A learned position-dependent strength $\pi_m$ and a context-relevance score $\kappa_m$ were then computed as

$$\pi_m = \sigma\left(a_{r_m}\right), \tag{9}$$

$$\kappa_m = \sigma\left(\text{MLP}_{\text{rel}}\left[\mathbf{h}_m; \mathbf{e}(r_m); \mathbf{h}_0; \mathcal{H}_m\right]\right), \tag{10}$$

where $a_{r_m}$ is a learned scalar associated with relative position $r_m$, $\mathbf{e}(r_m) \in \mathbb{R}^{H_{\text{ctx}}}$ is its learned positional embedding and $[\cdot\,;\cdot]$ denotes vector concatenation. The relevance prior was defined as

$$\alpha_m = \pi_m \kappa_m. \tag{11}$$

Thus, a genomic-context protein received a large prior only when both its relative genomic position and its predicted functional profile indicated relevance to the target protein.

After BiLSTM encoding, the target latent embedding served as the query, whereas the latent embeddings of the valid genomic-context proteins served as the keys and values for attention pooling. Let $\widetilde{\mathbf{h}}_0$ and $\widetilde{\mathbf{h}}_m$ denote the target and genomic-context latent embeddings, respectively. The pooling weights and decoded context embedding ($\mathbf{h_{ctx}}$) were

$$\beta_m = \underset{m}{\text{softmax}}\left[\frac{\widetilde{\mathbf{h}}_m^{\top} \widetilde{\mathbf{h}}_0}{\sqrt{H_{\text{ctx}}}} + \log\left(max\left(\alpha_m, \epsilon\right)\right)\right], \tag{12}$$

$$\mathbf{h_{ctx}} = \text{MLP}\left(\text{LayerNorm}\left(\sum_{m=1}^{M} \beta_m \widetilde{\mathbf{h}}_m\right)\right). \tag{13}$$

A learned output matrix converted the decoded context embedding into context-correction logits $\boldsymbol{\Delta} \in \mathbb{R}^C$. A per-function gate $\mathbf{g} \in [0,1]^C$ was computed by an MLP, denoted as $G$. Its inputs comprised the target logits, the context-correction logits and a five-dimensional vector $\boldsymbol{\eta}$. This vector contained the mean Bernoulli entropy of the target function probabilities, the mean, maximum and minimum of the corresponding entropies across valid genomic-context proteins, and the summed relevance prior. Therefore, the gate was calculated as:

$$\mathbf{g} = \sigma(G[\mathbf{s}_0, \boldsymbol{\Delta}, \boldsymbol{\eta}])\,. \tag{14}$$

The final target function probability $\widehat{\mathbf{y}}$ was computed as:

$$\mathbf{s}^* = \mathbf{s}_0 + \mathbf{g} \odot \boldsymbol{\Delta}, \qquad \widehat{\mathbf{y}} = \sigma(\mathbf{s}^*)\,. \tag{15}$$

### 2 Supplementary Tables

| **Model** | **Version** |
|---|---|
| ESM-C | 600M |
| ESM-2 | 650M |
| ProtBert | 420M |
| Empathi | 1.0.6 |
| DHR | v2 |
| Foldseek | 718d42176d2f67d36a60866fedfb881f8d5a7ebf |
| HMMER | v3.4 |
| MMseqs2 | 0b27c9d7d7757f9530f2efab14d246d268849925 |
| VPF-PLM | v1.0 |
| BACPHLIP | 0.9.6 |

Supplementary Table 1: Software versions used in this study.

| Parameter | HMMER (`phmmer`) | MMseqs2 | Foldseek |
|---|---|---|---|
| Sensitivity (`-s`) | Default | 7.5 | 9.5 |
| Alignment type | Default | Default | Type 2: local 3Di+AA alignment |
| Exhaustive search | Default | Enabled | Dataset-dependent[a] |
| Maximum prefilter candidates (`--max-seqs`) | Not specified | Not specified | 1000 |
| Minimum bitscore | 0 | Not specified | Not specified |
| Minimum query coverage | 0 | Not specified | Not specified |
| Minimum target coverage | 0 | Not specified | Not specified |
| Minimum sequence identity | Not specified | Not specified | Not specified |
| Sequence/alignment length cutoff | Not specified | Not specified | Not specified |
| Search-space size (`-Z`) | Full reference sequence count | Not specified | Not specified |
| GPU mode | Not specified | Not specified | Enabled |

*Notes.* "Not specified" indicates that the scripts do not explicitly set the parameter; built-in tool defaults still apply.
[a]Foldseek exhaustive search is enabled for efam, and disabled by default for MGV and phage lifestyle benchmark.

Supplementary Table 2: Inference hyperparameters for HMMER, MMseqs2, and Foldseek.

### 3 Supplementary Figures

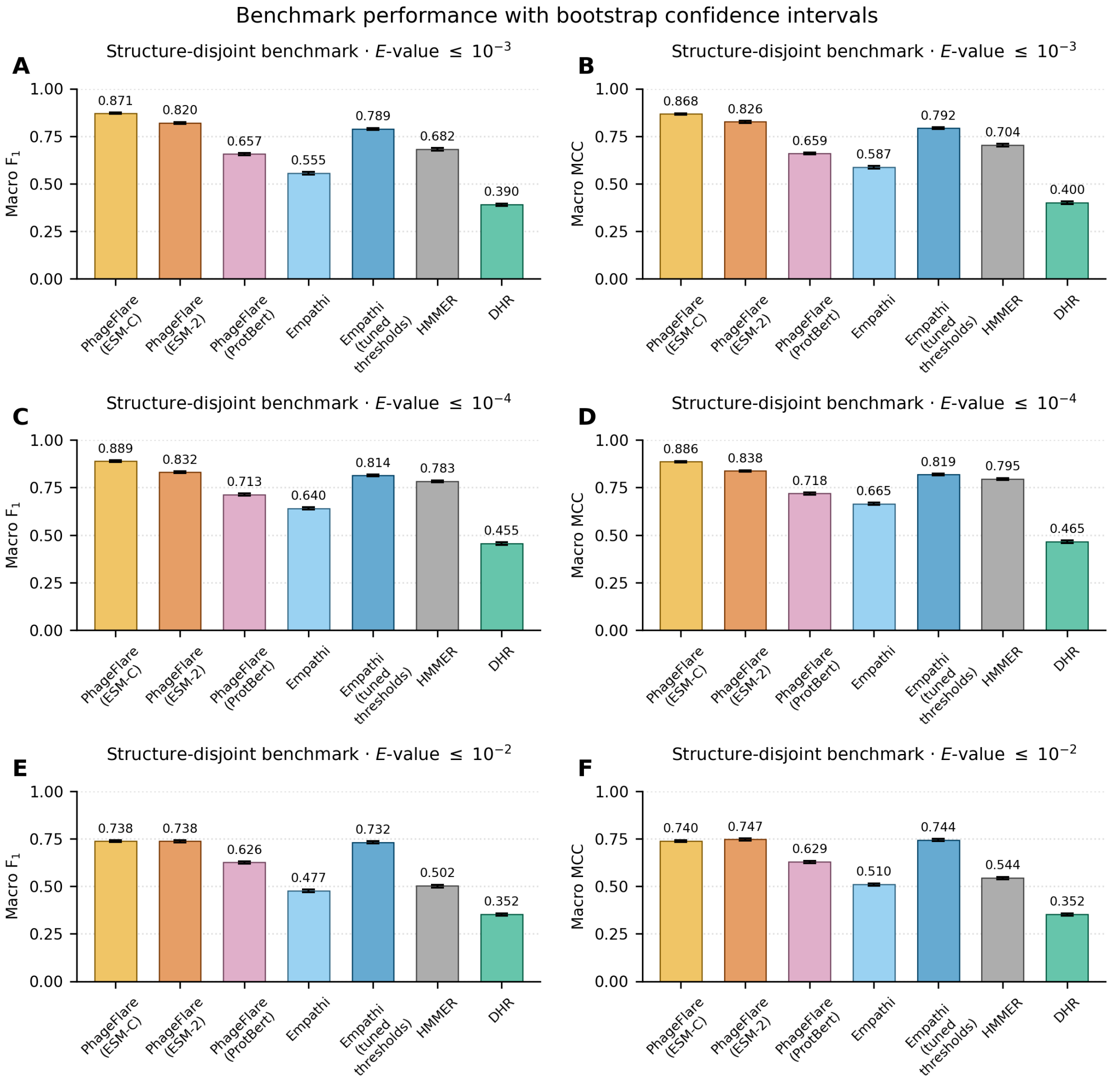


Supplementary Fig. 1: Bootstrap confidence intervals for benchmark performance across structure-disjoint test datasets. (**A**–**F**) Distributions of macro $F_1$ score (**A, C, E**) and macro MCC (**B, D, F**) on structure-disjoint test sets constructed under Foldseek $E$-value thresholds of $10^{-3}$ (**A, B**), $10^{-4}$ (**C, D**) and $10^{-2}$ (**E, F**). Bars display observed macro-averaged scores across 45 functional categories for PhageFlare (ESM-C), PhageFlare (ESM-2), PhageFlare (ProtBert), Empathi, Empathi (tuned thresholds), HMMER and DHR. Error bars denote pointwise 95% percentile confidence intervals derived from 10,000 paired protein-level bootstrap replicates. This figure reports confidence intervals for macro-$F_1$ and macro-MCC only. AUPRC values reported for HMMER and DHR in Fig. 2 were calculated from binary predictions because continuous scores were unavailable.

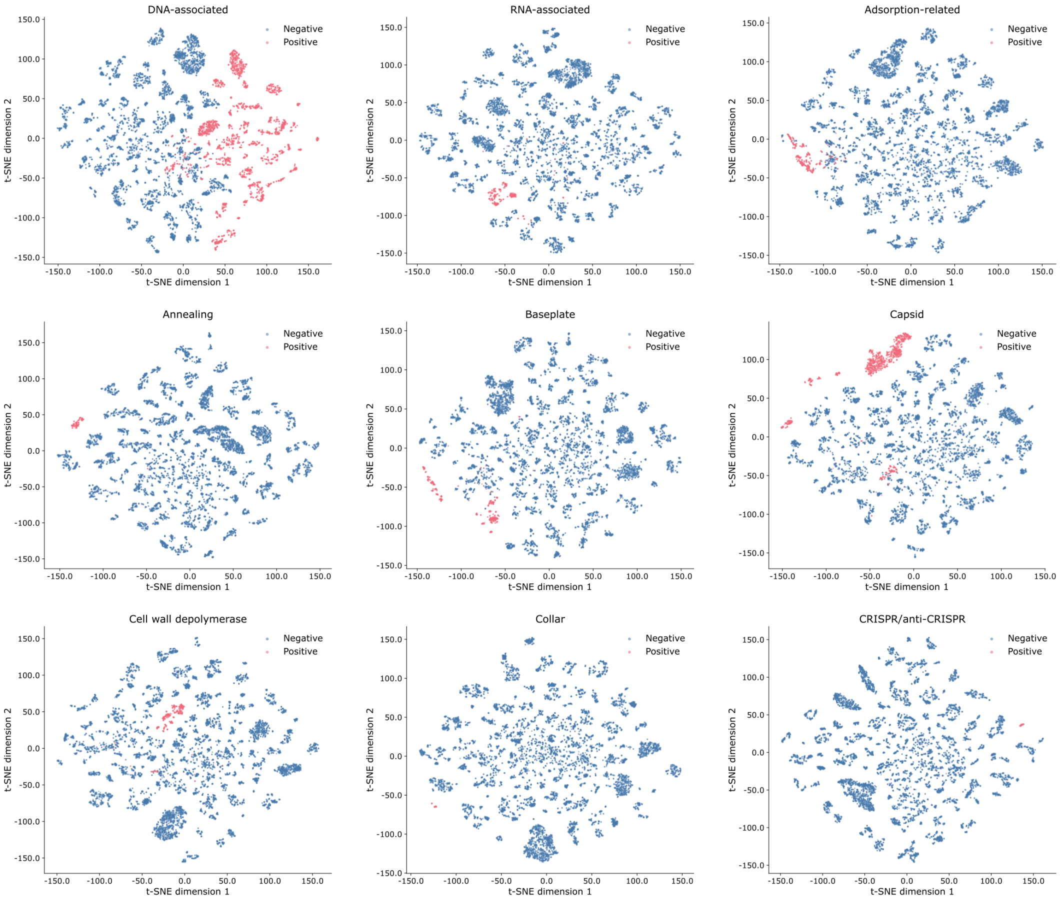


Supplementary Fig. 2: Two-dimensional t-SNE projections of target protein embeddings from the PhageFlare classifiers for DNA-associated, RNA-associated, adsorption-related, annealing, baseplate, capsid, cell wall depolymerase, collar, and CRISPR/anti-CRISPR.

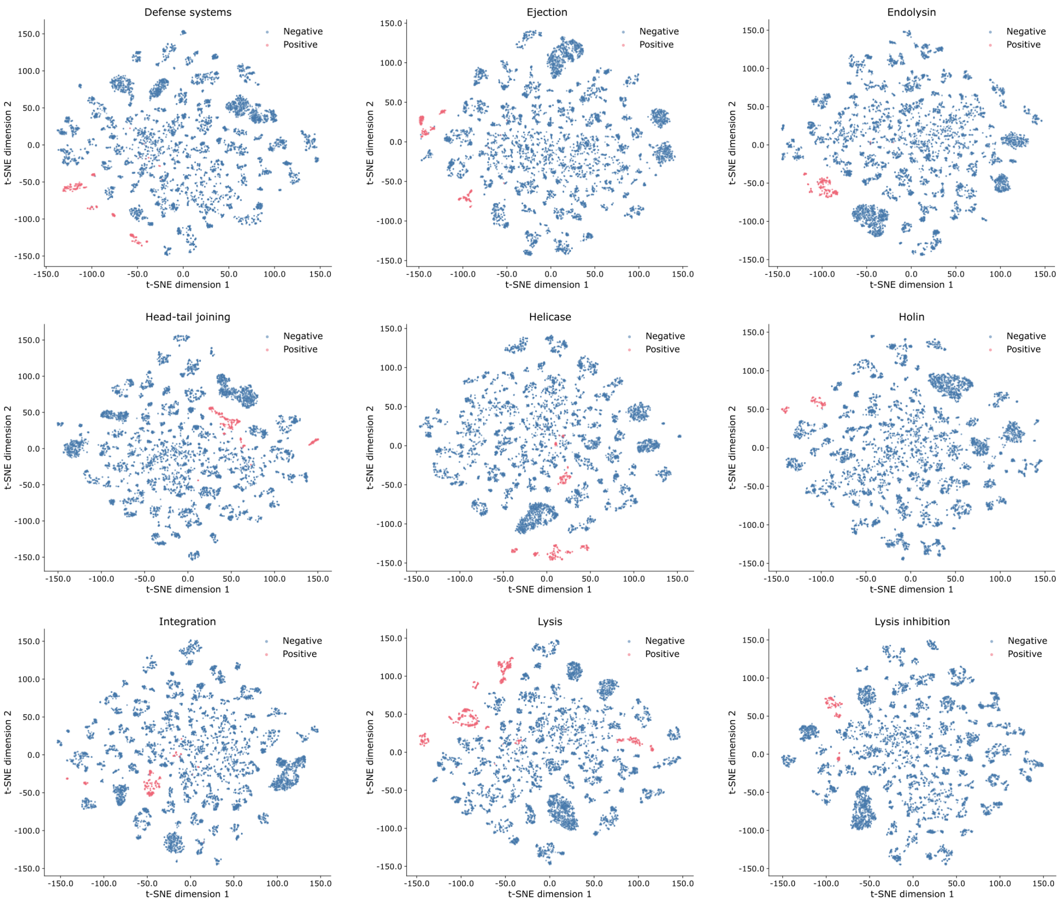


Supplementary Fig. 3: Two-dimensional t-SNE projections of target protein embeddings from the PhageFlare classifiers for defense systems, ejection, endolysin, head-tail joining, helicase, holin, integration, lysis, and lysis inhibition.

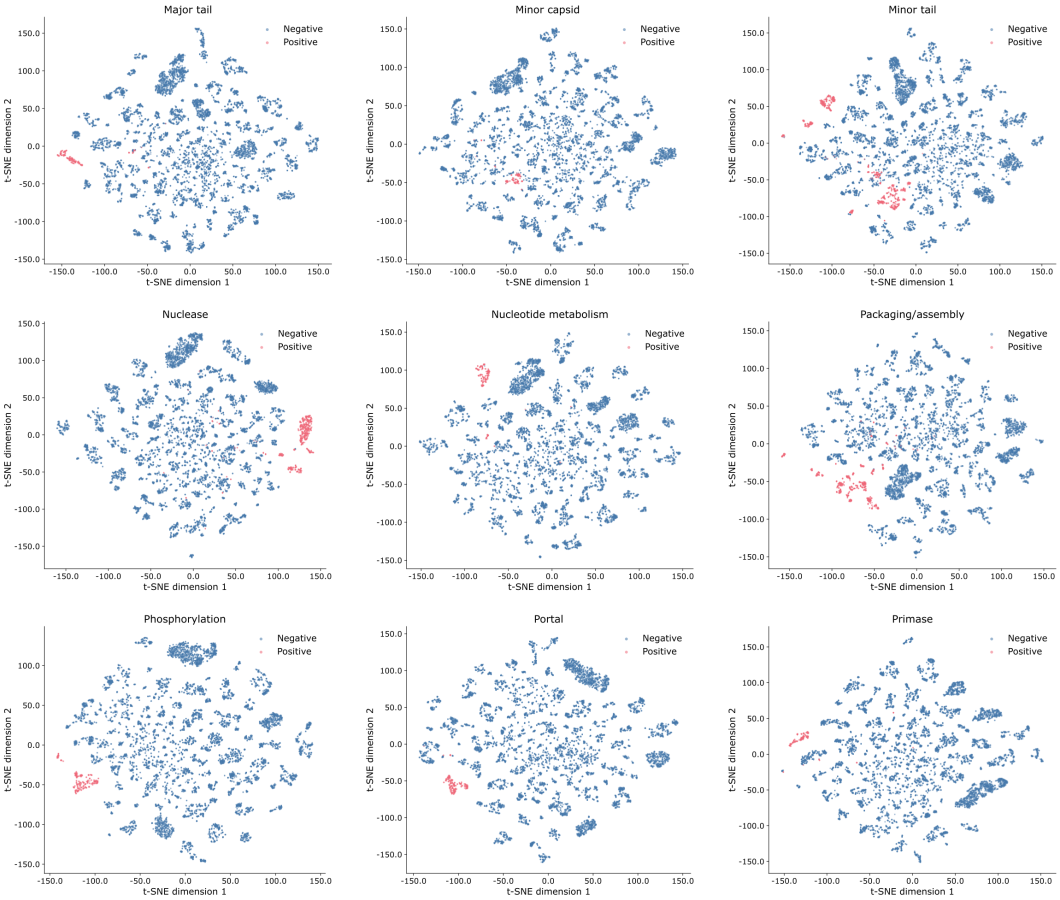


Supplementary Fig. 4: Two-dimensional t-SNE projections of target protein embeddings from the PhageFlare classifiers for major tail, minor capsid, minor tail, nuclease, nucleotide metabolism, packaging/assembly, phosphorylation, portal, and primase.

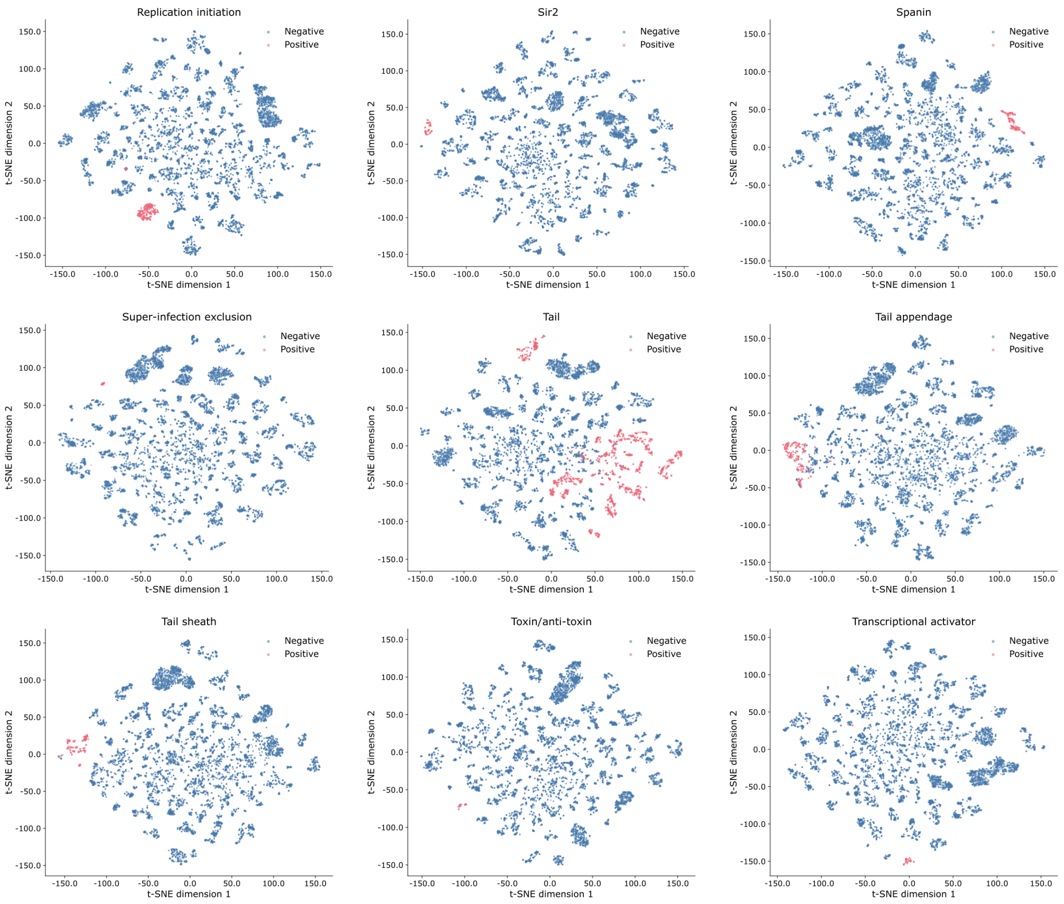


Supplementary Fig. 5: Two-dimensional t-SNE projections of target protein embeddings from the PhageFlare classifiers for replication initiation, Sir2, spanin, super-infection exclusion, tail, tail appendage, tail sheath, toxin/anti-toxin, and transcriptional activator.

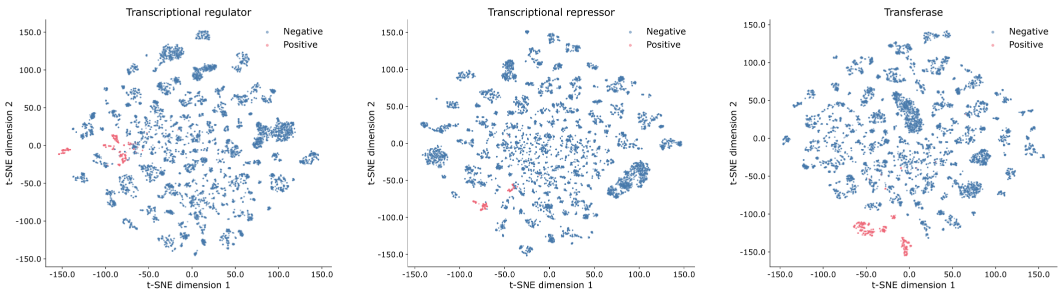


Supplementary Fig. 6: Two-dimensional t-SNE projections of target protein embeddings from the PhageFlare classifiers for transcriptional regulator, transcriptional repressor, and transferase.
